# Seq-Scope Protocol: Repurposing Illumina Sequencing Flow Cells for High-Resolution Spatial Transcriptomics

**DOI:** 10.1101/2024.03.29.587285

**Authors:** Yongsung Kim, Weiqiu Cheng, Chun-Seok Cho, Yongha Hwang, Yichen Si, Anna Park, Mitchell Schrank, Jer-En Hsu, Jingyue Xi, Myungjin Kim, Ellen Pedersen, Olivia I. Koues, Thomas Wilson, Goo Jun, Hyun Min Kang, Jun Hee Lee

## Abstract

Spatial transcriptomics (ST) technologies represent a significant advance in gene expression studies, aiming to profile the entire transcriptome from a single histological slide. These techniques are designed to overcome the constraints faced by traditional methods such as immunostaining and RNA *in situ* hybridization, which are capable of analyzing only a few target genes simultaneously. However, the application of ST in histopathological analysis is also limited by several factors, including low resolution, a limited range of genes, scalability issues, high cost, and the need for sophisticated equipment and complex methodologies. Seq-Scope—a recently developed novel technology—repurposes the Illumina sequencing platform for high-resolution, high-content spatial transcriptome analysis, thereby overcoming these limitations. Here we provide a detailed step-by-step protocol to implement Seq-Scope with an Illumina NovaSeq 6000 sequencing flow cell that allows for the profiling of multiple tissue sections in an area of 7 mm × 7 mm or larger. In addition to detailing how to prepare a frozen tissue section for both histological imaging and sequencing library preparation, we provide comprehensive instructions and a streamlined computational pipeline to integrate histological and transcriptomic data for high-resolution spatial analysis. This includes the use of conventional software tools for single cell and spatial analysis, as well as our recently developed segmentation-free method for analyzing spatial data at submicrometer resolution. Given its adaptability across various biological tissues, Seq-Scope establishes itself as an invaluable tool for researchers in molecular biology and histology.

**KEY POINTS:**

- The protocol outlines a method for repurposing an Illumina NovaSeq 6000 flow cell as a spatial transcriptomics array, enabling the generation of high-resolution spatial datasets.
- The protocol introduces a streamlined data analysis pipeline that produces a spatial digital gene expression matrix suitable for various single-cell and spatial transcriptome analysis methods.
- The protocol allows for the capture of histology images from the same tissue section subjected to spatial transcriptomics analysis and allows users to precisely align the transcriptome dataset with the histological image using fiducial marks engraved on the flow cell surface.
- Leveraging commonly available Illumina equipment, the protocol offers researchers ultra-high submicrometer resolution in spatial transcriptomics analysis with a comprehensive pipeline, rapid turnaround, cost efficiency, and versatility.

## INTRODUCTION

### Development of the protocol

Conventional histology, employing methods such as immunohistochemistry and RNA *in situ* hybridization, is limited to detecting only a single or a few specific molecular entities on a tissue slide. As a result, the information acquired from one experimental run is restricted, likely missing crucial biological details.

To address these challenges, the emerging field of spatial transcriptomics (ST) aims to analyze the expression of the entire transcribed genome from a single histological slide. However, existing ST technologies face significant challenges due to their low resolution or limited capacity for high-throughput analysis. For example, the widely used Visium platform from 10x Genomics offers a resolution of 100 μm^1^, which falls below the 40 µm resolution discernible by the naked human eye. Although newer methods employing microbead arrays or microfluidics have enhanced resolution to approximately 10–20 µm^2–4^, they have yet to match the precision of traditional optical microscopy. High-resolution techniques, such as sequential *in situ* hybridization and *in situ* sequencing, are available in commercial ST platforms, including 10x Xenium^5^, Nanostring CosMx^6^, and Vizgen MERSCOPE^7^. However, these approaches involve extensive staining/imaging processes and require intensive computational image analysis, limiting the number of genes that can be analyzed simultaneously within a practical time frame and budget. Such constraints on the number of target genes per project likely restrict research focus and introduce bias, potentially impeding access to crucial biological insights that would otherwise be accessible with a more comprehensive approach.

Single cell RNA-seq (scRNA-seq) is an unbiased approach that comprehensively examines transcriptomic heterogeneity across individual cells^8^. However, due to the nature of tissue dissociation and single cell isolation, all current scRNA-seq methods have clear limitations. First, scRNA-seq omits many labile cell types that lyse during tissue dissociation and cell sorting. Second, extensive tissue dissociation and cell sorting procedures cause physical and chemical stresses that may elicit stress responses in cells. Large and lipid-laden cell types such as myofibers and adipocytes are not amenable for cell sorting by microfluidics or microwells. Finally, and most obviously, scRNA-seq eliminates the spatial information of single cells in the histological space, resulting in loss of important biological information, such as tissue zonation niches and cell-cell interactions.

Therefore, improving the resolution, throughput, and scalability of ST is crucial for overcoming the limitations of currently available ST and scRNA-seq technologies. Recently, we developed Seq-Scope^9^—which represents the first published high-resolution ST method capable of providing submicrometer resolution (0.5–0.7 µm on average), while monitoring the whole transcriptome at the sequence level. This enables single-cell and even subcellular analysis and represents a dramatic improvement in spatial resolution. For instance, the spatial throughput (spatial units per area) of Seq-Scope is over 10,000 times higher than conventional 10x Visium (with a 100 µm resolution) and 4–8 times higher than the recently released 10x Visium HD platform (released in January 2024, with a 2 µm resolution^10^; peer-reviewed publication is not yet available).

Seq-Scope utilizes the Illumina sequencing platform, repurposing its flow cells into high-resolution spatially barcoded arrays. Publication of our method^9^ was followed by several papers employing similar approaches utilizing independent sequencing platforms, such as DNA Nanoball (DNB) sequencing^11^ and a custom polony sequencing platform^12^ that can also provide resolutions similar to Seq-Scope. However, Seq-Scope’s unique characteristics and capabilities, outlined below, make it especially useful and accessible for investigators seeking to utilize high-resolution spatial techniques.

The Seq-Scope protocol was developed by building upon existing scRNA-seq and ST technologies. The protocol incorporates well-established components from PCR-based scRNA-seq library creation, as referenced in the Drop-Seq^13^ and BD Rhapsody^14^ protocols, along with general tissue handling techniques from ST protocols described in the original spatial transcriptomics methodology^15^ and the commercially available 10x Visium kit protocols^16, 17^. It also utilizes the STARsolo scRNA-seq alignment tool^18^ for data processing. These established methods, combined with our innovative adaptation of the Illumina sequencing flow cell for ST, have led to the creation of a new experimental and computational workflow, as outlined in our previous publications on the Seq-Scope methodology^9^ and the STtools computational analysis package^19^. Our method is compatible with most Illumina sequencing platforms, including MiSeq^9^ and HiSeq^20–22^, and has been independently adapted for 3-dimensional analysis on the NovaSeq^23^, demonstrating its wide applicability and reproducibility. We further optimized the method for a 7 mm × 7 mm imaging area on the NovaSeq 6000 flow cell, chosen due to its suitability for the majority of histological applications and comparability to the 6.5 mm × 6.5 mm area used by 10x Visium. Our updated protocol successfully reproduced the major findings detailed in our original Seq-Scope study^9^, as outlined below and in **Anticipated Results**.

### Application of the method

Seq-Scope has proven to be highly effective for profiling the spatial transcriptome of gastrointestinal organs such as the liver and colon^9^. It has also been instrumental in revealing the intricate details of the human skin spatial transcriptome^20^. Our extensive in-house analyses further demonstrate that Seq-Scope is versatile and compatible with a wide range of tissue types, including those that pose challenges in single-cell or spatial analyses such as skeletal muscle with elongated and multinucleated myofibers^22^.

### Comparison with other methods

Advantages

1. **Submicrometer Resolution:** Seq-Scope sets a new benchmark in spatial transcriptomics resolution. Offering a center-to-center resolution of 0.5–0.7 µm, it stands as the first published technique with submicrometer resolution for whole transcriptome analysis, providing the highest resolution among sequence-based ST technologies. This high resolution enables Seq-Scope to offer detailed insights into the spatial organization of tissues at both the single-cell and subcellular levels. For instance, its subcellular analysis capability, demonstrated by comparing nuclear and cytoplasmic spatial transcriptomes^9^, successfully replicated findings obtained from physical fractionation^24^.
2. **High Capture Efficiency:** Seq-Scope demonstrated a remarkable capture capacity, achieving over 23 UMI/µm² in a colon dataset^9^, making it one of the leading technologies in terms of capture efficiency in the ST field^25^. Its demonstrated capacity is surpassed only by the recently released Decoder-Seq^26^ and Patho-DBiT^27^, which offer 40–60 UMI/µm² but at a lower resolution of approximately 10–100 µm. This high capture capacity, coupled with submicrometer spatial resolution, ensures comprehensive and dense sampling of transcripts, resulting in a detailed representation of the spatial transcriptome. To achieve this, Seq-Scope utilizes a straightforward chemistry that boosts transcriptome capture efficiency. Such simplicity increases reliability and extends Seq-Scope’s accessibility to a broader range of researchers. Despite variations in capture rates across tissue sections due to differences in RNA concentration (∼6.7 UMI/µm² in the liver dataset^9^, influenced by its large vasculature/sinusoidal volume, and ∼1.7 UMI/µm² in the skin dataset^20^, affected by the skin’s dense extracellular matrix layer that is rich in non-cellular components and devoid of RNA), Seq-Scope’s high capture efficiency ensures that maximum information is extracted from any given sample.
3. **Leveraging the Illumina Sequencing Platform:** Seq-Scope benefits from the high fidelity and robustness of the widely used Illumina sequencing platform. This integration ensures reliable and accurate sequencing results—essential for detailed transcriptomic analysis—and offers precision over other methods that rely on a DNB-seq or custom sequencing array. Furthermore, given the ubiquity of Illumina sequencers, most research groups or institutions can replicate this protocol without the need to purchase new equipment. Moreover, the procedures for generating flow cell arrays can be readily performed by most sequencing service providers across the world. This increases accessibility and convenience for researchers, ensuring that Seq-Scope remains a viable option for a wide range of laboratories, regardless of their equipment inventory.
4. **Accessibility of Materials:** Beyond the Illumina sequencing platform, all of the reagents and consumables required for Seq-Scope are commercially available, significantly easing the adoption of this method in various research settings. The widespread availability of these materials ensures that a multitude of researchers can employ this advanced technology without substantial logistical challenges.
5. **Precise Annotation of Histopathology:** Unlike certain methods (e.g., Stereo-seq using opaque DNB-seq flow cells), the Illumina sequencing flow cell is transparent, making Seq-Scope compatible with most histological methods including hematoxylin and eosin (H&E) staining. This compatibility facilitates the identification of tissue histopathology and its relationship with the spatial transcriptome. In addition, the alignment of histology with spatial transcriptomes can be precisely achieved using preexisting fiducial marks engraved on the flow cell, which in turn enables accurate transcriptomic annotation of tissue histopathology.
6. **Streamlined Computational Analysis:** Seq-Scope data analysis is supported by a streamlined computational package that simplifies data processing and produces a spatial digital gene expression matrix useful for various single cell and spatial analysis methods. In addition, we provide example scripts for identifying spatial factors using Latent Dirichlet allocation (LDA) or Seurat multidimensional clustering^28^, followed by projection of these factors into histological space at submicrometer resolution using FICTURE, our recently developed algorithm^29^.
7. **Rapid Turnaround and Scalability:** Seq-Scope is structured around two sequencing steps (1st-Seq and 2nd-Seq), which are efficiently executed on the established Illumina system. Users concentrate primarily on tissue processing and library preparation, tasks that are typically completed within two days, with an additional day for Chip processing. Moreover, the protocol’s design allows a single technician to simultaneously process over 10 samples in a single batch, demonstrating the method’s exceptional scalability.
8. **Cost-Effectiveness:** Using the same method published previously^9^, we scaled up the volumes and optimized the steps to use the NovaSeq 6000 flow cell, which provides a larger analysis area while reducing sequencing costs. For a single imaging area of 7 mm × 7 mm, the array synthesis cost is approximately $150, with minimal reagent costs approaching $50. The total cost varies by sequencing depth, but even with a modest sequencing budget of ∼$250, the spatial hepatocellular transcriptome was comprehensively profiled, comparable to what we demonstrated in our initial dataset^9^. In contrast, the 10x Visium array—despite offering a much lower resolution—incurs substantially higher costs. Processing four 6.5 mm × 6.5 mm imaging areas with Visium costs approximately $5,000—averaging $1,250 per area and excluding the additional recommended sequencing cost of approximately $1,200 per area. 10x Visium HD, though still offering lower resolution, is anticipated to be pricier than its predecessor. This comparison highlights the advantage of Seq-Scope in delivering high-resolution ST at substantially lower cost, establishing it as a cost-effective choice for laboratories—particularly those operating on tight budgets.
9. **Versatility and Potential for Future Development:** The protocol provides detailed steps on how to use an Illumina sequencer to produce a high-resolution spatial array. It allows individual researchers to create custom arrays by applying minor modifications to adapter sequences, potentially enabling additional spatial applications such as spatial epigenome or antibody tag sequencing. The transparency and flexibility inherent to 1^st^-Seq array synthesis in conjunction with its numerous applications signify that Seq-Scope is not merely a current solution for transcriptomics but also provides a platform for future innovations. Its potential for multi-omics analysis and other applications opens new avenues in ST research.

Disadvantages

1. **Relatively Lower Resolution Compared to Imaging-based Technologies:** Although Seq-Scope boasts the highest resolution among spatially barcoded whole transcriptome sequencing technologies, its resolution does not theoretically surpass the best resolution of the optical microscope. For instance, its resolution is comparable to a microscope with 10X objectives (∼0.7 µm) but lower than that with 100X objectives (∼0.2 µm). Therefore, current resolution of Seq-Scope is lower than targeted transcriptomics techniques using high-magnification imaging methods such as *in situ* hybridization or *in situ* sequencing technologies.
2. **Limited Coverage of Rarely Expressed Transcripts:** Given that Seq-Scope captures the entire transcriptome in an unbiased manner, just like bulk RNA-seq and other single-cell and spatial RNA-seq methods, its transcriptome representation is proportional to the abundance of individual genes. Therefore, rarely expressed genes may have limited coverage compared to methods that target specific genes. While deeper sequencing of the library can mitigate this issue in principle, targeted transcriptomics techniques based on *in situ* imaging may be more suitable when the primary focus is on monitoring some rarely expressed target molecules rather than on having transcriptome-wide coverage.

## Experimental Design

The Seq-Scope procedure is divided into two separate sequencing steps. The first array sequencing (1st-Seq) begins as conventional Illumina sequencing of a single-stranded oligonucleotide library encoding a high-definition map coordinate identifier (HDMI) as a random sequence that is attached to PCR adapters and an oligo-dT RNA capture probe. After 1st-Seq, the sequenced flow cell is retrieved and disassembled to expose the surface covered with amplified DNA clusters containing HDMI sequences. The output of 1st-Seq is a data table that associates all HDMI barcode sequences from the physical array along with their corresponding coordinate information.

Next, the 1st-Seq HDMI array is biochemically treated to enable poly-A capture, which is achieved by exposing its 3’ oligo-dT sequences. Thereafter, the target tissue is mounted onto the 1st-Seq HDMI array and briefly fixed in place. Conventional imaging, such as H&E histology staining or fluorescence imaging, can be performed at this stage. Subsequently, RNAs are released by partial proteolytic digestion of tissues using collagenase and pepsin and captured onto the 1st-Seq HDMI array. The tissue-array sandwich then undergoes reverse transcription, labeling the HDMI-oligo-dT molecule with cDNA sequences. Through subsequent secondary strand synthesis, alkaline elution, and PCR amplification steps, the chimeric molecular library of HDMI-cDNA is prepared as an Illumina-compatible library that is then sequenced on an Illumina sequencer. This encompasses 2nd-Seq, the spatial profiling of the transcriptome in the tissue. The output of the 2nd-Seq data analysis is another data table with two paired-end reads—one end with HDMI information and the other end with cDNA sequence information.

For each HDMI sequence, 1st-Seq generates spatial coordinate information while 2nd-Seq captures cDNA information. We construct a spatial gene expression matrix by integrating the data from both sequencing steps, which then serves as the basis for various analyses. This integration is facilitated by a computational pipeline we developed^19^ that processes the datasets for compatibility with conventional single-cell optimized software packages such as Seurat^28^ and Scanpy^30^. In addition, our recently developed tool—FICTURE—allows for pixel-level inference of individual transcriptome phenotypes^29^, effectively leveraging the high-resolution spatial information inherent to the Seq-Scope dataset.

## Expertise Needed to Implement the Protocol

Array and library sequencing must be performed by technicians with experience running next-generation sequencing on Illumina equipment. This can be accommodated by most sequencing cores and companies with minor modifications to their workflow as described in the protocol provided below. Given that the sequencing procedures will likely be outsourced, the current protocol would be widely accessible without the need to own or operate specialized equipment.

Tissue processing and library preparation can be performed by any laboratory staff trained in basic molecular biology and histology procedures. The data analysis methods demonstrated in this protocol can also be performed by anyone who can install and run standardized pipelines for processing next-generation sequencing results on a Unix-based system.

## Limitations

The current protocol is limited to detecting the transcriptome; expanding it to include other modalities such as proteomes, genomes, and epigenomes would necessitate modifications. These expansions in applications could be achieved by adapting published methods, originally designed for lower-resolution techniques^31–35^, for use with the high-resolution Seq-Scope array described in this protocol.

In addition, the current protocol employs poly-A capture using oligo-dT probes and a random priming method, which sacrifices information in the upstream sequences. Compared to the full-length capture methods previously used to interrogate transcriptomes for ST^15^, this method offers greater sensitivity as it can capture damaged transcripts that cannot be reverse transcribed in full length. However, it loses 5’ sequences, which often contain important information, such as B-cell and T-cell receptor sequences^36–38^. Modifying the current protocol would be necessary to enable the capture of such sequences.

Finally, successful implementation of Seq-Scope is contingent upon the preparation of high-quality fresh frozen sections. The current protocol has not been validated with sections that have compromised RNA quality, nor with fixed frozen sections or formalin-fixed paraffin-embedded (FFPE) tissues—all of which pose known challenges for ST studies. Addressing these challenges may involve modifications to the workflow to better accommodate FFPE tissues. This could include enhancing transcript recovery methods^39, 40^, labeling fragmented transcripts with poly-A tails^27^, and/or employing indirect transcriptome capture techniques using *in situ* hybridized probes that can anneal to fragmented mRNA sequences^41^.

## MATERIALS

### Biological Materials

Adult mouse liver tissue was previously collected from 8-week-old male C57BL/6 wild-type mice immediately after CO_2_ euthanasia. The tissue was embedded in an optical cutting temperature (OCT) compound on cooled isopentane (2-methylbutane) in a liquid nitrogen bath and stored at –80 °C.

CRITICAL: If the sample is obtained externally, the freezing procedure should be conducted immediately after tissue collection as described in Step 2-1 (Tissue Preparation) or by using a similar method. Delays in this procedure could compromise the RNA quality. Inappropriate freezing conditions may lead to the formation of ice crystals, causing freezing tissue damage. Although tissues snap-frozen in liquid nitrogen sometimes work well for small samples, they often suffer from damage due to uneven freezing conditions caused by bubbles.

### Oligonucleotides

Oligonucleotides can be ordered from major vendors including IDT (Integrated DNA Technologies, Inc., Coralville, IA, USA) and Eurofins Genomics (Eurofins Genomics LLC, Louisville, KY, USA).

- HDMI32-DraI: CAAGCAGAAGACGGCATACGAGATTCTTTCCCTACACGACGCTCTTCCGATCTNNVNBV NNVNNVNNVNNVNNVNNVNNVNNNNNTCTTGTGACTACAGCACCCTCGACTCTCGC TTTTTTTTTTTTTTTTTTTTTTTTTTTTTTAAAGACTTTCACCAGTCCATGATGTGTAGATCT CGGTGGTCGCCGTATCATT
- Read1-DraI: ATCATGGACTGGTGAAAGTCTTTAAAAAAAAAAAAAAAAAAAAAAAAAAAAAAGCGAGAGTCGAGGGTGCTGTAGTCACAAGA

CRITICAL: Given the unusual length of these oligonucleotides, they must be synthesized as Ultramer DNA oligos by IDT (or as EXTREmer at Eurofins). All other oligos can be synthesized as conventional DNA oligos. Based on our experience, PAGE purification is not necessary.

- Randomer v2: TCAGACGTGTGCTCTTCCGATCTNNNNNNNNB
- T-Stopper TTTTTTTTTTTTTTTTTTTTTTTTTTTTTT/3Phos/
- RPEPCR*F: TCTTTCCCTACACGACGC*T*C
- RPEPCR*R: TCAGACGTGTGCTCTTCC*G*A
- WTA1*F: AATGATACGGCGACCACCGAGATCTACACTCTTTCCCTACACGACGCTCT*T*C
- WTA1*R*N705: CAAGCAGAAGACGGCATACGAGAT<u>AGGAGTCC</u>GTGACTGGAGTTCAGACGTGTGCTCTT CC*G*A
  ○ The underlined index sequences are modifiable and can be substituted with alternative indices to create new indexing primers such as:
    ∎ WTA1*R*N706: replace underlined nucleotides with CATGCCTA
    ∎ WTA1*R*N707: replace underlined nucleotides with GTAGAGAG
    ∎ WTA1*R*N708: replace underlined nucleotides with CCTCTCTG

### Oligonucleotide Setup

- The oligonucleotides are reconstituted in UltraPure water at 100 µM concentration, aliquoted, and stored at –20 °C.

### Reagents

- 0.1 N Hydrochloric acid (EMD Millipore/MilliporeSigma, Burlington, MA, USA; cat. no. 1.09060.1000)
- 2-Mercaptoethanol (Sigma-Aldrich, St. Louis, MO, USA; cat no. M3148)

CAUTION: Exercise caution while handling 2-mercaptoethanol due to its potent smell and ability to irritate the skin, eyes, and respiratory system. Use only in a well-ventilated area or under a fume hood and wear appropriate protective equipment such as gloves and goggles.

- Acetic acid, glacial (Fisher Chemical/Thermo Fisher Scientific, Waltham, MA, USA; cat. no. A38-212)

CAUTION: Acetic acid is highly corrosive and can cause severe burns to skin and eyes, as well as respiratory irritation. Use with appropriate protective equipment and in a well-ventilated area or fume hood.

- Agarose (Fisher Bioreagents/Thermo Fisher Scientific, cat. no. 9012-36-6)
- AMPure XP purification beads (Beckman Coulter, Brea, CA, USA; cat. no. A63881)
- Collagenase I (Thermo Fisher, cat. no. 17018-029)
- Dako Bluing Buffer (Dako/Agilent Technologies, Santa Clara, CA, USA; cat. no. CS702)
- Dako Mayer’s Hematoxylin (Dako/Agilent, cat. no. S3309)
- DNA Clean & Concentrator®-25 (Zymo Research, Irvine, CA, USA; cat. no. D4014)
- dNTPs (New England Biolabs, Ipswich, MA, USA; cat. no. N0477L)
- DPBS (Dulbecco’s phosphate buffered saline; Gibco/Thermo Fisher, cat. no. 14190-144)
- Dra I restriction enzyme (New England Biolabs, cat. no. R0129)
- 0.5 M EDTA pH 8.0 (Invitrogen™/Thermo Fisher, Carlsbad, CA, USA; cat. no. AM9261)
- Eosin Y solution (Sigma-Aldrich, cat. no. HT110216)
- Ethyl alcohol (Sigma-Aldrich, cat no. E7023)
- Exonuclease I (New England Biolabs, cat. no. M2903)
- Exonuclease I Reaction Buffer (New England Biolabs, cat. no. B0293)
- Ficoll® PM 400 (Sigma-Aldrich, cat. no. F4375-10G)
- Gel Loading Dye, Purple (6X; New England Biolabs, cat. no. B7024)
- Glass beads (BioSpec, Bartlesville, OK, USA; cat. no. 11079127)
- Glycerol (Fisher Bioreagents/Thermo Fisher, cat. no. 56-81-5)
- HBSS (Hank’s Balanced Salt Solution; Gibco/Thermo Fisher, cat. no. 14175-095)
- Isopentane (2-methyl butane; EMD Millipore/MilliporeSigma, cat. no. MX0760-1)
- Isopropanol (Sigma-Aldrich, cat. no. 19516)

CAUTION: Exercise careful handling of isopentane and isopropanol to minimize the risk of chemical exposure and flammability hazards.

- Kapa HiFi Hotstart ReadyMix (KAPA Biosystems, Inc./Roche, Wilmington. MA, USA; cat. no. KK2602)
- Klenow Fragment (exonuclease-deficient; New England Biolabs, cat. no. M0212)
- Liquid nitrogen and dry ice

CAUTION: Use liquid nitrogen and dry ice with care to avoid frostbite injuries.

- Maxima 5X RT buffer (Thermo Fisher, cat. no. EP0751)
- Maxima H Minus Reverse Transcriptase (Thermo Fisher, cat. no. EP0751)
- NEBuffer 2 (New England Biolabs, cat. no. B7002)
- Neutralization Buffer (Zymo Research, cat. no. D4027-3-50)
- O.C.T. Compound Clear (Fisher Healthcare/Thermo Fisher, cat. no. 4585)
- Paraformaldehyde solution, 16% (Electron Microscopy Sciences, Hatfield, PA, USA; cat. no. 15170)

CAUTION: Handle paraformaldehyde with extreme caution due to its potential for toxic exposure and irritation to skin, eyes, and respiratory system.

- Pepsin (Sigma-Aldrich, cat. no. P7000)
- Proteinase K (New England Biolabs, cat. no. P8107)
- Qubit™ dsDNA High-Sensitivity assay kit (Invitrogen™/Thermo Fisher, cat. no. Q32854)
- rCutSmart™ Buffer (New England Biolabs, cat. no. B6004)
- RNase Inhibitor [Lucigen Biosearch Technologies (LGC), Middleton, WI, USA; cat. no. 30281]
- RNeasy Plus Mini Kit (Qiagen, Hilden, Germany; cat. no. 74134)
- Sodium chloride (Sigma-Aldrich, cat. no. S3014)
- Sodium n-dodecyl sulfate (SDS; EMD Millipore/MilliporeSigma, cat. no. 428015)
- SYBR™ Gold Nucleic Acid Gel Stain (10,000X Concentrate in DMSO; Invitrogen/Thermo Fisher, cat. no. S11494)
- Tris Base, ULTROL® Grade (EMD Millipore/MilliporeSigma, cat. no. 648313)
- UltraPure distilled water (Invitrogen™/Thermo Fisher, cat. no. 10977-015)
- Zymoclean™ Gel DNA Recovery Kit (Zymo Research, cat. no. D4001)

### Supplies

- Custom Frame Adapter (University of Michigan Duderstadt Center Fabrication Facility, Ann Arbor, MI, USA)
  ○ Printed using Stratasys J850™ using default white material.
  ○ A 3D model file (in STL) is provided in Supplementary Data 1 with which the part can be fabricated by most 3D printing service providers.
- Custom Silicone Isolator (Grace Bio-Labs, Bend, OR, USA; cat. no. JTR25-A-1.0, RD501346)
  ○ Sketch drawing with specifications (in PDF) is provided in Supplementary Data 2.
  ○ With this information, Grace Bio-Labs can fabricate custom parts with the custom specifications.
- DNA LoBind™ Tubes, 1.5 mL (Eppendorf, Hamburg, Germany; cat.no. 0030 108.051)

CRITICAL: Use LoBind tubes to prevent DNA libraries from adhering to the tube walls. This adherence can reduce yield and lower concentration, an issue commonly encountered with conventional tubes.

- Latex gloves
- MB22 Premier Microtome Blade (Epredia, Kalamazoo, MI, USA; cat. no. 3050822)
- Metal beaker
- Micropipette filter tips
- Micropipettes (Conventional set: 1000 µL, 200 µL, 20 µL, 2 µL)
- Microscope cover glass (Mercedes Scientific, Lakewood Ranch, FL, USA; cat. no. #MER R2575)
- NovaSeq S4 35 cycle reagent kit (Illumina, San Diego, CA, USA)
- PCR tubes (ProAmp™, Alkali Scientific, Fort Lauderdale, FL, USA; cat. no. PC7064)
- Qubit assay tubes (Invitrogen™/Thermo Fisher, cat. no. Q32856)
- Shandon™ Base Mold 15 × 15 mm (Thermo Scientific/Thermo Fisher, cat. no. 41741)
- Styrofoam box

### Equipment

- 37 °C Incubator (New Brunswick™ Galaxy™ 170S; New Brunswick Scientific, Edison, NJ, USA)
- 42 °C Incubator (Boekel Scientific™ model no. Rapid FISH; Feasterville-Trevose, PA, USA)
- Agarose gel electrophoresis system
- Benchtop centrifuge for 1.5 mL microcentrifuge tubes (Eppendorf, model no. 5424)
- Agilent 2100 BioAnalyzer (Agilent)
- Chemical fume hood
- Cryostat (Epredia™, model no. Cryostar™ NX50)
- Illumina NovaSeq 6000 system (Illumina)
- LabChip (Revvity, Waltham, MA, USA)
- Light microscope with built-in camera and bright-field channel (any; Keyence, Osaka, Japan; model no. BZ-X810)
- Magnetic stand for 1.5 mL tubes (Invitrogen/Thermo Fisher, model no. DynaMag™-2)
- NanoDrop™ Spectrophotometer (Thermo Fisher)
- PCR system (Applied Biosystems/Thermo Fisher, model no. ProFlex)
- Precision Glass Cutter (UTILE, model no. FU-200)
- Qubit™ 3 Fluorometer (Invitrogen/Thermo Fisher, cat. no. 33216)
- Razor blade
- Scalpel (Dynarex Medi-Cut, Montvale, NJ, USA; cat. no. 4110)
- Scriber/Etching Pen (IMT, cat. no. 8800)
- TapeStation (Agilent).
- ThermoMixer® (Eppendorf, model no. F1.5)

### Reagent Setup

- 0.45 M Tris-acetate pH 6.0 buffer: weigh 5.4 g Tris base powder and dissolve in 90 mL UltraPure water. Adjust the pH to 6.0 with acetic acid and fill to 100 mL with UltraPure water. Store at room temperature.
- 4% Paraformaldehyde: prepare by adding 1 mL 16% paraformaldehyde to 3 mL PBS. Store at 4 °C.

CRITICAL: For optimal results, make 4% paraformaldehyde fresh prior to use; 16% paraformaldehyde stock can be stored for a week after opening the ampule.

- Collagenase I stock solution (50 U/μL): prepare by dissolving collagenase powder in HBSS buffer to achieve a concentration of 50 U/μL. Subsequently, pass the solution through a 0.45-µm filter. Aliquot and store at –20 °C
- Pepsin stock solution (0.1g/mL): Measure 0.1 g of pepsin power, and dissolve in 1 mL of UltraPure water. Mix well by pipetting. Aliquot and store at –20 °C

CRITICAL: Do not vortex collagenase I and pepsin stock solutions. Aliquot the stock solutions into small volumes to avoid multiple freeze-thaw cycles as they substantially decrease the enzymatic activities of these stock solutions.

### System Requirements for Computational Analysis

- A high-performance computing system with 4 or more CPUs with 64GB or more memory, with >1TB disk. A larger number of CPUs (>10) and memory (>300GB) is preferred for deeply sequenced data.
- Compatible with UNIX-based operating systems (e.g., Ubuntu, CentOS, MacOS).
- Compatible with a standalone machine or HPC with SLURM workload manager.

## PROCEDURE

### A. Experimental Procedures

The current experimental procedures are divided into three major steps: Step 1 produces a spatially barcoded array, Step 2 attaches the tissue to the array, performs tissue imaging, and prepares the tissue for library construction, and Step 3 captures the tissue transcriptome through reverse transcription and processes the spatially barcoded cDNA into a next-generation sequencing library (Fig. 1).

**Fig. 1.**
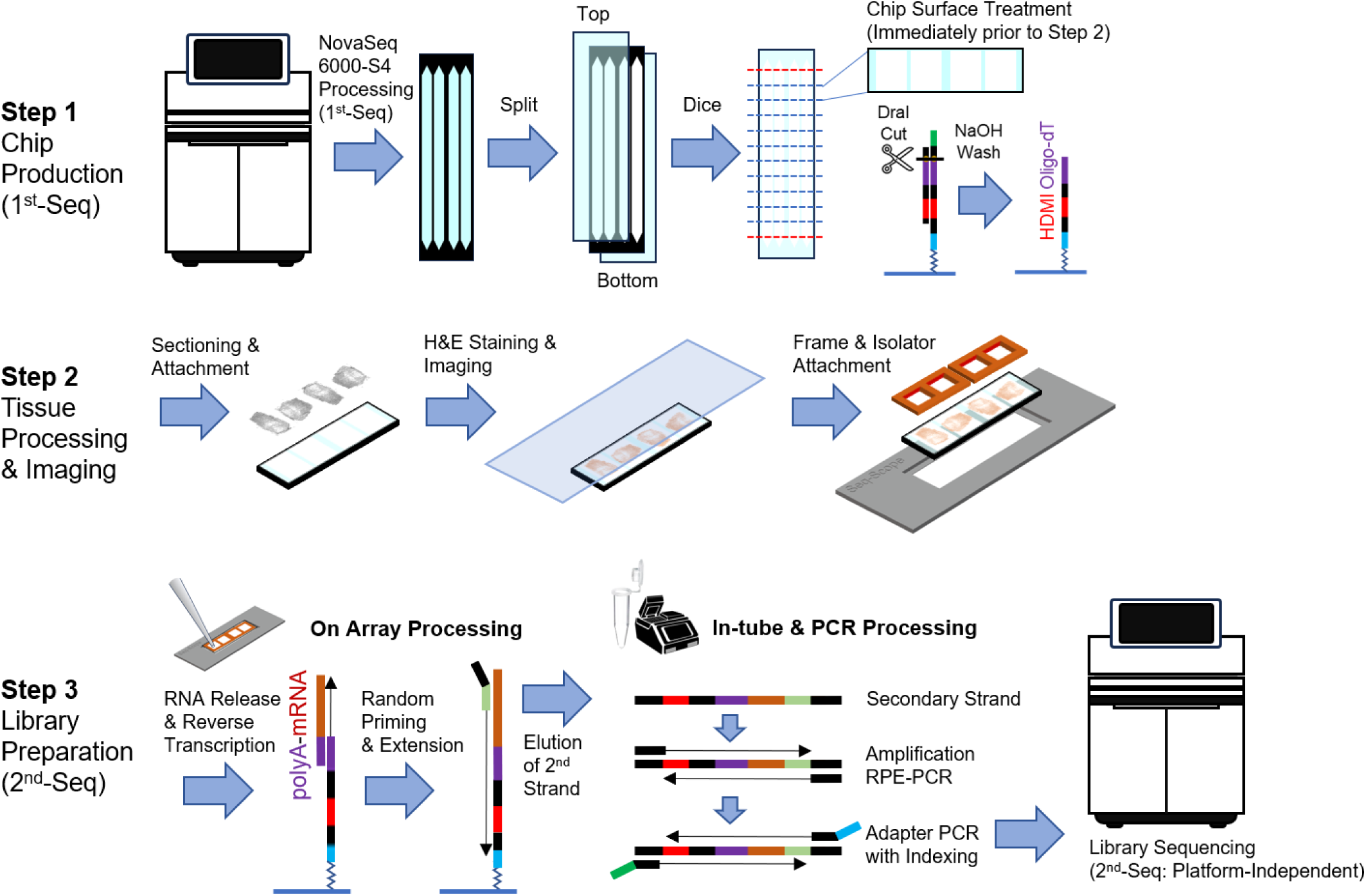
Overview of Experimental Procedures.

#### STEP 1. Seq-Scope Chip Production (1^st^-Seq)

##### 1-1) NovaSeq 6000 Operation: TIMING ∼15 hours (instrument can run overnight)

Consult the Illumina® (San Diego, CA, USA) NovaSeq 6000 Sequencing System Guide (https://support.illumina.com/content/dam/illumina-support/documents/documentation/system_documentation/novaseq/1000000019358_17_novaseq-6000-system-guide.pdf) for instructions on how to thaw and prepare the NovaSeq 6000 S4 v1.5 (35 cycle) reagent kit, applying the following modifications as described in the Critical Steps below:

1. HDMI32-DraI (ssDNA oligonucleotide) is the input library. Read1-DraI is the custom Read 1 sequencing primer. CRITICAL STEP: The input library is an ssDNA oligonucleotide that is not amenable to typical concentration/sizing methods such as TapeStation. Moreover, qPCR quantification is inaccurate because this library contains highly conserved sequences. We also found that the molar concentration provided by IDT is not a reliable indicator of the library’s clustering efficiency. It is important to note that libraries ordered from IDT with identical sequences vary significantly in clustering efficiency. Therefore, Illumina’s recommended library loading concentrations cannot be determined using the standard quantitation methods mentioned above. We instead recommend that this critical parameter be determined empirically through serial dilution and titration, as outlined below.
2. Initially, the XP workflow described in the sequencer guide is used to test four different concentrations in each of the four lanes of the S4 flow cell in the first round of titration:
  a. Lane 1: 0.5 nM
  b. Lane 2: 1.0 nM
  c. Lane 3: 1.5 nM
  d. Lane 4: 2.0 nM CRITICAL STEP: Optimal loading concentration varies widely between oligonucleotide batches and should be determined via titration experiments on each batch. Seq-Scope is robust across a range of cluster densities, with both underclustered (0.5 nM) and overclustered (2.0 nM) Seq-Scope arrays generating useful ST data; however, the number of usable pixels and overall data yield will be lower than that generated using optimized concentrations—determined to be 1.5 nM in our case.
3. The instrument is configured to perform only Read 1 cycles for sequencing by synthesis (SBS), with no Index 1, Index 2, or Read 2 steps so that cluster re-synthesis does not occur.
4. Prior to sequencing, the instrument operator must replace the bleach-wash solution in well #17 of the cluster cartridge with laboratory-grade water to ensure that the array clusters do not deteriorate during the unavoidable post-run wash performed automatically upon completion of sequencing. This will preserve cluster integrity for subsequent procedures.
5. Once sequencing is complete, the instrument operator can store the flow cell at 4 °C for later delivery to the investigator. The operator should then perform a maintenance wash as soon as possible to adequately remove template residue from the instrument’s fluidic components. Meanwhile, the researcher can access the run statistics using Illumina’s Sequence Analysis Viewer (SAV; https://support.illumina.com/downloads/sequencing-analysis-viewer-3-0-product-documentation.html). Fig. 2 shows run stats retrieved from SAV for the titration run detailed in Step 2 above. The base composition at each cycle should reflect the unique pattern of HDMI32 (NNNNNBNNBNNBNNBNNBNNBNNBNNBVNBNNA, the reverse complement of HDMI sequences in HDMI32-DraI; Fig. 2a).
6. When the procedure is performed by outside sequencing cores and centers, the investigator should request delivery of all files, including thumbnails and run statistics. As accessed via SAV, the run statistics is especially helpful for inspecting array quality. Nevertheless, the Read 1 FASTQ file is sufficient for bioinformatic analysis in that it contains all necessary information, including HDMI sequences, lane and tile information, as well as XY coordinates.
7. Inspect cluster density. It is typical to achieve > 2 million PF clusters per mm^2^. In the example shown in Fig. 2b and 2c, Lane 1 is slightly underclustered, while Lanes 3 and 4 are slightly overclustered. Although the number of PF clusters is highest in Lane 2, our bioinformatic assessment indicated that slight overclustering is preferable in that it inhibits cluster duplication. Consequently, the experiment detailed in Fig. 2 points to an optimal loading concentration of 1.5 nM. Once optimized, the standard onboard loading workflow should be used in place of the XP workflow, reducing costs and improving overall sequencing quality and number of PF clusters. Another round of optimization may be necessary upon switching to onboard clustering, as this workflow optimizes enzyme activity. Illumina recommends 30% higher loading concentrations for on-board loading compared to the XP workflow; however—given the more robust enzyme activity—this increase must be evaluated to prevent potential overloading.

**Fig. 2.**
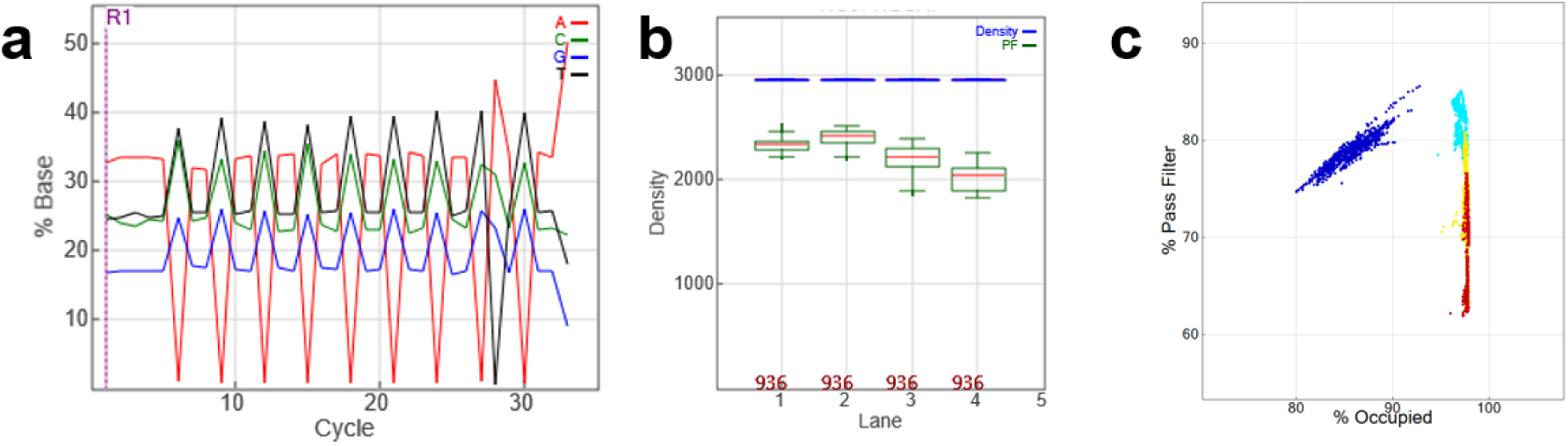
Example outputs from the Illumina SAV application showing. (a) base composition at each cycle, (b) raw (Density) and pass filter (PF) cluster density in each lane, and (c) scatterplot of cluster occupancy (x axis) and % PF in each lane (blue – Lane 1, cyan – Lane 2, yellow – Lane 3, red – Lane 4). Each dot represents an individual tile in one of the four flow cell lanes.

CRITICAL STEP: We observed that repeated freeze-thaw cycles negatively impact the cluster efficiency of the HDMI32-DraI library. Therefore, HDMI32-DraI stock solutions should be aliquoted and frozen to minimize freeze-thaw damage.

PAUSE POINT: The sequenced flow cell can be stored at 4°C for up to 2 months.

##### 1-2) Flow Cell Disassembly and Dicing: TIMING ∼2 hours

1. Ensure you have a clean, well-lit workspace and gather all necessary tools, including a scalpel and dicing station [Precision Glass Cutter (UTILE, model no. FU-200)] with a straight edge and two magnet holders.
2. Carefully use the scalpel to separate the flow cell into its three main components: the top flow cell layer, the middle plastic spacer, and the bottom flow cell layer. Refer to Fig. 3a for guidance and consult Video 1 for a demonstration.
3. Examine the top and bottom flow cell layers for patterned areas where clusters are formed. These areas can be identified by a characteristic rainbow reflection, as shown in Fig. 3b.
4. Horizontally flip the top layer of the flow cell so that the side with the clusters is facing upward. The bottom layer can remain in its original position.
5. Move to the dicing station. Follow the procedure demonstrated in Fig. 3c and Video 2 and outlined in Fig. 3d to prepare the glass for dicing.
6. Using a straightedge and two magnet holders, secure the flow cell in its dicing position.
7. Carefully score along the predetermined line. Ensure that the line is accurately scored for a clean break.
8. Gently snap the glass along the scored line to break it into pieces (see Video 2).
9. You will end up with a total of 22 diced glass fragments, designated B01–B11 and T01–T11.
10. Remove and discard fragments B01, B11, T01, and T11, as the clusters in these areas were not sequenced in the NovaSeq 6000 operation.
11. You should now have 18 viable diced glass fragments, numbered B02–B10 and T02–T10. Each of these fragments contains 4 lanes, labeled A–D or 1–4, resulting in 72 distinct imaging areas. Note that each flow cell fragment measures 1.1 cm × 4 cm and has four imaging areas, each with a maximum imaging space of approximately 7 mm × 11 mm. Each fragment is now referred to as a Seq-Scope Chip, or simply “Chip.”
12. Chips can be dried and stored at 4℃ for up to two months without substantial loss of functionality.

CAUTION: Handle with care during the glass dicing and chip generation process, as the edges of the cut glass can be sharp.

**Fig. 3.**
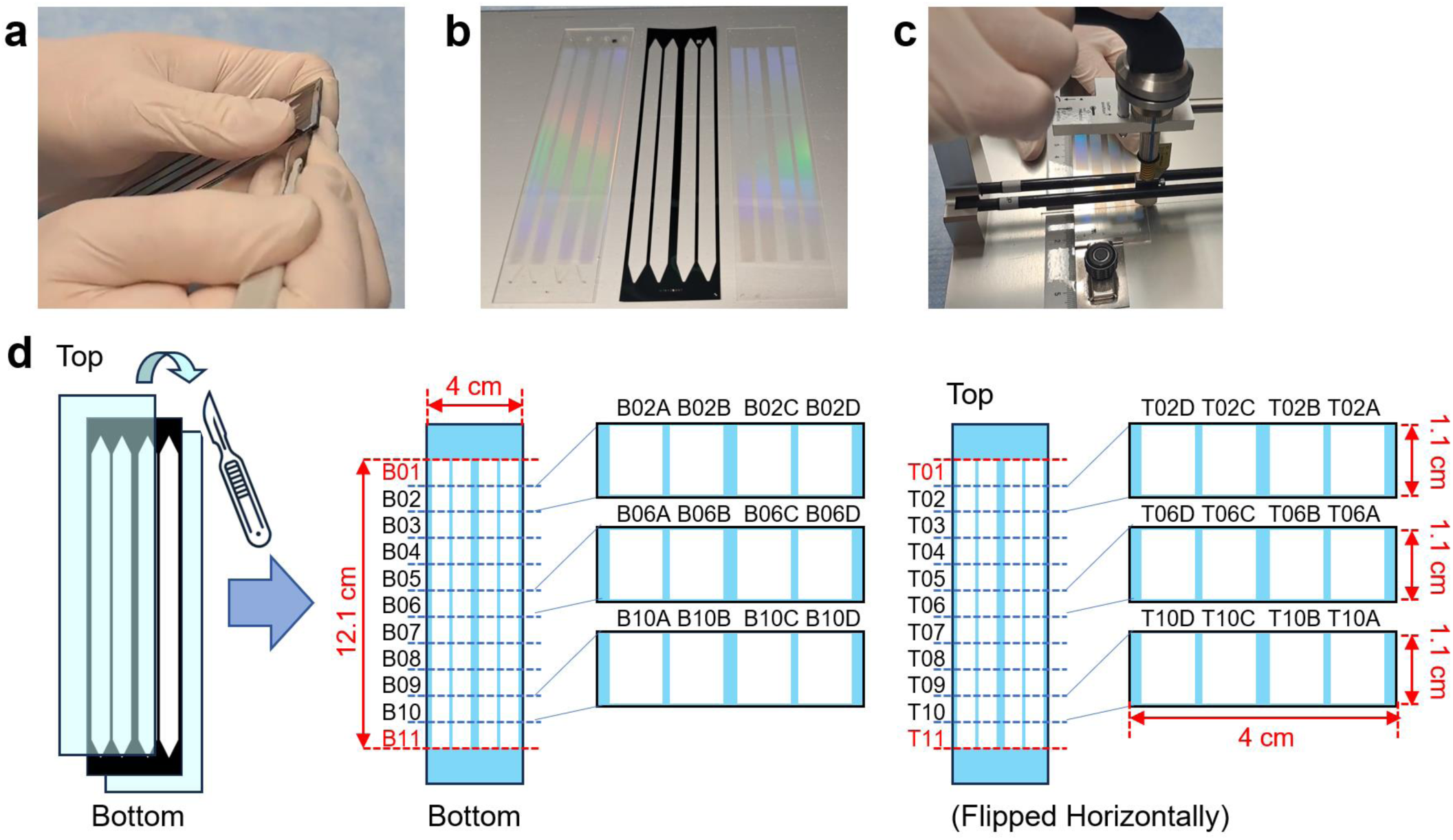
Flow cell disassembly and dicing. (a) Disassembly of NovaSeq S4 flow cell using a scalpel. (b) Flow cell disassembled into top, middle, and bottom layers. The top layer is upside down to position the cluster surface upward. (c) Glass dicing procedure. (d) Schematic of the flow cell diced into separate fragments and imaging areas.

CRITICAL: Ensure that the working area and dicing station are thoroughly cleaned; the handler must wear fresh protective latex or nitrile gloves to avoid contaminating the chip surface. In general, the handler should wear protective gloves throughout the procedure.

PAUSE POINT: The diced chip can be stored at 4°C for up to 2 months.

##### 1-3) Seq-Scope Chip Surface Treatment: TIMING ∼16 hours (including an overnight step)

1. Retrieve the Chip from the refrigerator and warm it to room temperature.
2. Set up the DraI digestion solution using the following formulation. 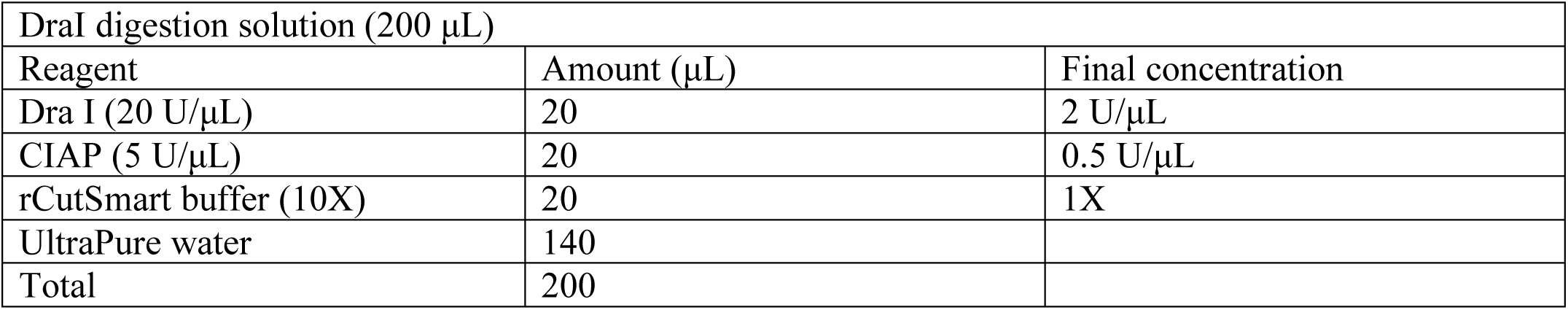
3. Begin with a 100 mm Petri dish to create a humidified chamber (see Fig. 4a).
4. Place a Kimwipe tissue at the bottom of the dish and saturate it thoroughly with water to ensure adequate humidity.
5. Lay a piece of Parafilm over the moist Kimwipe, stretching it out to cover the surface without tearing.
6. On top of the Parafilm, carefully dispense drops of DraI digestion solution as depicted in Fig. 4a, ensuring they are well spaced.
7. Gently place the Chip upside down onto the drops of DraI solution as depicted in Fig. 4b. This orientation allows the clusters on the Chip to be exposed directly to the enzyme for processing.
8. Seal the Petri dish securely to maintain the chamber’s humidity, ensuring that no air can escape and disrupt the humid environment.
9. Place the sealed humidified chamber into an incubator set at 37 ℃ for overnight incubation. This step is crucial for the enzymatic digestion process, allowing the DraI enzyme to effectively process the clusters on the Chip.
10. After the overnight incubation, retrieve the humidifying chamber from the incubator.
11. Carefully retrieve the Chip and wash it with excess water by placing it into a Petri dish containing UltraPure water. Repeat three times.
12. Dry the Chip using a Kimwipe without touching the cluster surface.
13. Return to the humidifying chamber and prepare it again with fresh Parafilm.
14. Set up the Exonuclease I solution using the following formulation. 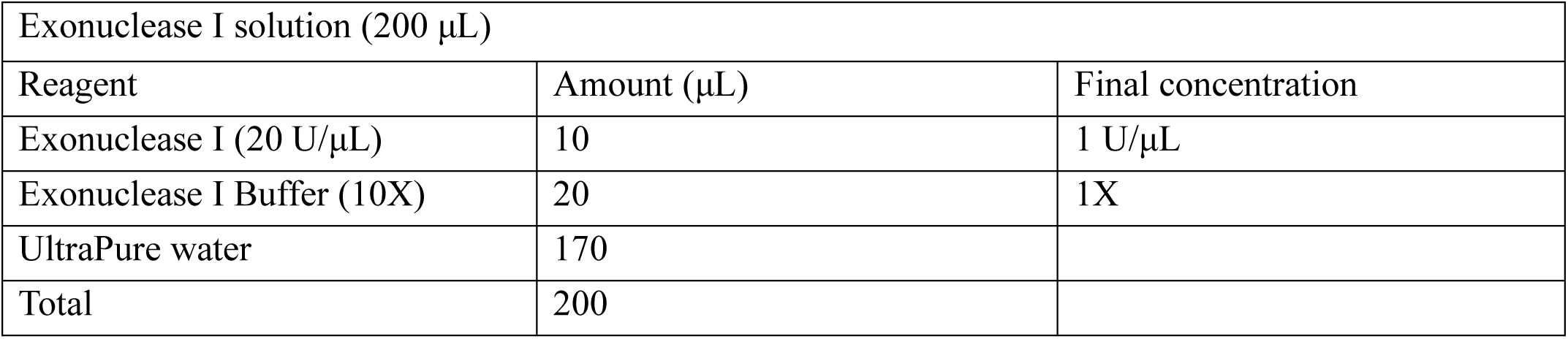
15. Carefully dispense drops of Exonuclease I solution on the Parafilm as depicted in Fig. 4a, ensuring that they are well spaced.
16. Gently place the Chip upside down onto the drops of Exonuclease I solution as depicted in Fig. 4b. This orientation allows the clusters on the Chip to be exposed directly to the enzyme for processing.
17. Seal the Petri dish securely to maintain the chamber’s humidity, ensuring that no air can escape and disrupt the humid environment.
18. Insert the sealed humidified chamber into an incubator preset at 37 °C for 45 minutes. This essential step is intended to remove the P5 lawn along with any non-specific single-stranded DNA. During this process, the HDMI-containing molecules on the Chip remain safeguarded as they continue to form duplexes with Read1-DraI.
19. After the incubation is complete, retrieve the humidifying chamber from the incubator.
20. Carefully retrieve the Chip and wash it with excess water by placing it into a Petri dish containing UltraPure water. Repeat the washing three times and dry using a Kimwipe without touching the cluster surface.
21. Carefully dispense drops of 0.1N NaOH solution on the Parafilm and place the Chip upside down onto the drops of NaOH solution. Incubate at room temperature for 5 minutes. Repeat the procedure three times.
22. Carefully retrieve the Chip and wash it with excess 0.1 M Tris pH 7.5 by placing it into a Petri dish containing 0.1 M Tris pH 7.5. Repeat the washing three times.
23. Carefully retrieve the Chip and wash it with excess Ultrapure water by dropping it into a Petri dish containing the water. Repeat the washing three times.
24. The Chip is now ready for tissue attachment and can be stored at 4 ℃.

CRITICAL STEP: It has been observed that surface-treated Chips are stable for approximately one week. Beyond this period—particularly after a couple of weeks—there may be a decline in the quality of the libraries obtained from Chips that are stored long-term. Based on these occasional outcomes, it is recommended that Chips be stored in an unprocessed state for long-term preservation. Chip processing should ideally occur immediately prior to the actual experiment and library construction. This approach helps ensure the integrity of the Chip and the quality of the library, maximizing the potential for successful outcomes.

**Fig. 4.**
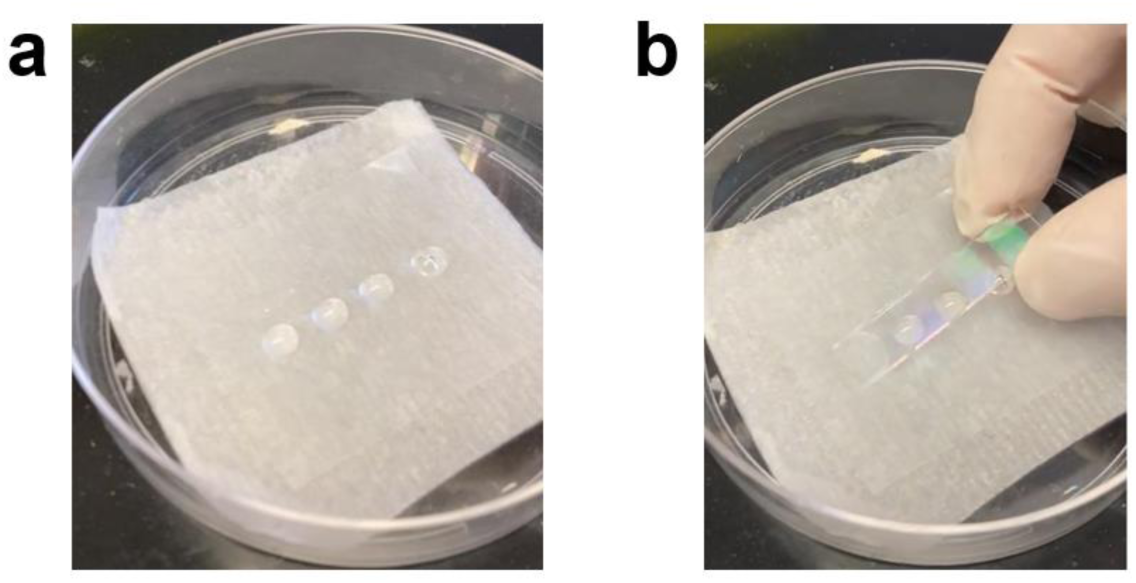
Liquid Handling in Seq-Scope Chip Surface Treatment. (a) Enzymatic and chemical solutions were first placed on Parafilm in a humidified chamber. (b) The chip is then overlaid onto the droplet solutions to treat the cluster surface.

PAUSE POINT: The surface-treated chip can be stored at 4°C for up to 1 week.

#### STEP 2. Tissue Preparation, Attachment, and Imaging

##### 2-1) Tissue Preparation (Freezing Instruction after Collection): TIMING ∼1 hour

1. Prepare an isopentane (2-methylbutane) and liquid nitrogen (LN_2_) bath.
  1) Fill a metal beaker to half capacity with isopentane, ensuring adequate volume to completely immerse the tissue. Position the beaker inside a liquid nitrogen (LN_2_) bath, aligning the liquid levels of both substances for optimal contact (refer to Fig. 5a).
  2) Incubate for 15 minutes. Cool in LN_2_ until the isopentane becomes viscous, indicated by a white film that covers the bottom of the beaker.
  3) Extended exposure to LN_2_ may cause the isopentane to solidify. Should this happen, momentarily remove the beaker from the bath to allow the isopentane to liquefy again before re-immersing it. CRITICAL STEP: It is crucial that both the isopentane and its container are thoroughly cooled to match the temperature of the LN_2_, thereby achieving a stable condition signaled by the cessation of boiling by the LN_2_. Initiating tissue immersion prematurely into a container that may not have reached the requisite low temperature can potentially cause the formation of ice crystals in the tissue that adversely affect its histological integrity.
2. Gently envelop fresh tissue samples in room temperature OCT compound inside a Petri dish. Ensure that the tissues are segmented into small portions; for example, mouse liver was segmented into 5 mm^3^ cubes.
3. Use a spatula to gently transfer the OCT-encapsulated tissue into a cryomold appropriate for the tissue size.
4. Ensure that the tissue is fully immersed in OCT to achieve complete coverage. If necessary, mark the cryomold to indicate the tissue’s orientation.
5. Carefully immerse the cryomold containing the tissue into the isopentane using forceps—avoiding complete submersion—as illustrated in Fig. 5b.
6. Maintain the position of the cryomold in the isopentane until the OCT becomes firm and turns opaque white; this requires approximately 30 seconds for liver tissue. CRITICAL STEP: The freezing duration must be determined empirically and should be brief. If the tissue is left in the isopentane bath for a prolonged period, freezing artifacts such as tissue distortion appear more frequently.
8. After the tissue has completely frozen, transfer the cryomold onto dry ice.
9. For prolonged preservation, store the cryo-embedded tissue in an airtight container at –80 °C. PAUSE POINT: The frozen sample can be stored at –80°C for up to 6 months.

**Fig. 5.**
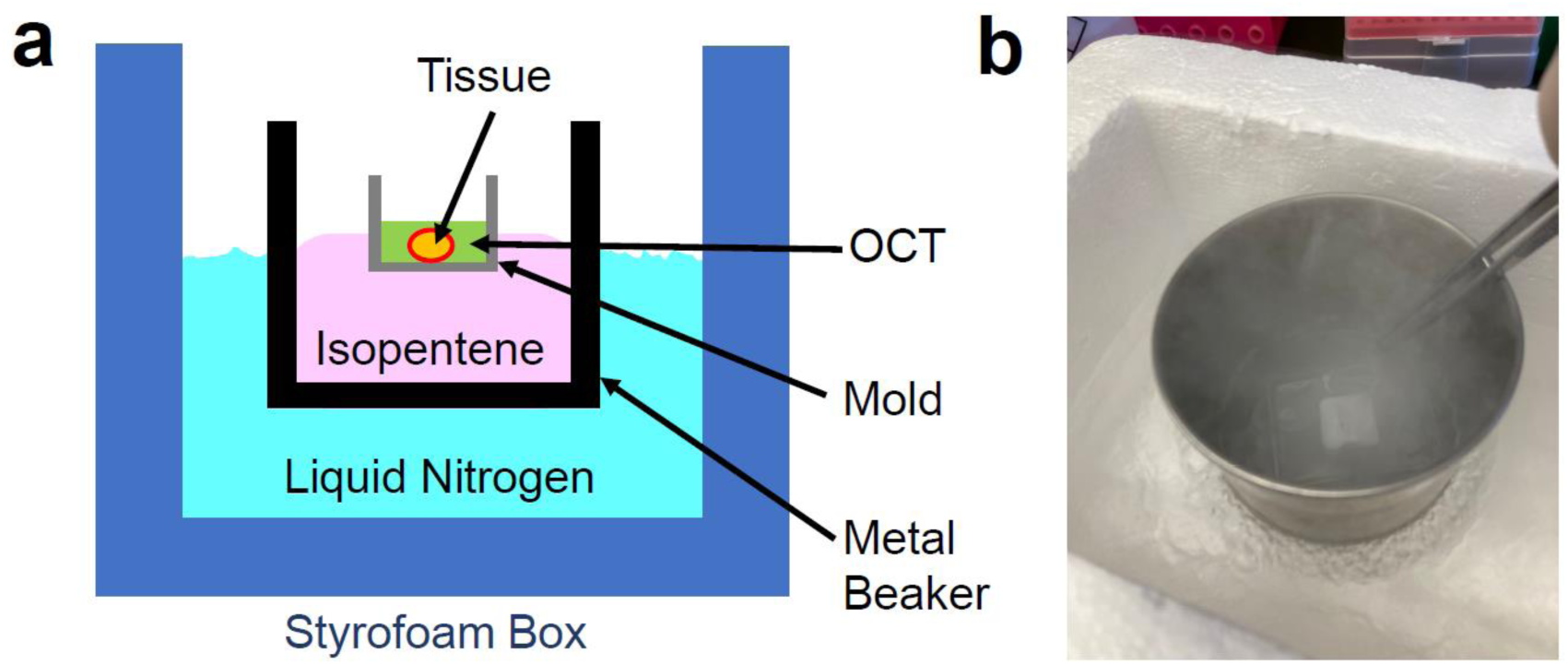
Tissue Freezing Chamber. (a) Schematic instructions for preparing a tissue freezing chamber. An LN_2_-chilled isopentane bath provides a medium for rapidly freezing OCT tissue block. (b) Demonstration of the tissue freezing procedure.

##### 2-2) Tissue sectioning and attachment: TIMING ∼2 hours

1. Chill the microcentrifuge tube, razor, brush, and forceps in the cryostat chamber at –20 °C to avoid thawing the tissue sections.
2. Retrieve the fresh-frozen tissue embedded in OCT from the –80 °C freezer. Place it in dry ice for > 30 minutes, then in a –20 °C cryostat for > 20 minutes. CRITICAL STEP: If you are unsure of histological tissue quality or RNA integrity (e.g., old tissue blocks or tissue blocks obtained outside the lab), you may choose to perform the quality check procedures described in the next section. If so, proceed to 2-3) Quality check for tissue histology and RNA integrity.
3. Section the tissue with a cutting angle of 5 degrees, ensuring each slice is 10 μm thick (refer to Fig. 6a).
4. Carefully place the Chip over the cryostat-sectioned tissue slices, ensuring that the cluster surface is facing downward, allowing the sections to adhere precisely to the proper surface areas of the Chip (refer to Fig. 6b).
5. Once attached, flip the Chip and gently press your finger against its underside to melt the tissue onto its surface (refer to Fig. 6c).
6. Repeat the tissue attachment procedure until tissue sections fully cover all four units of the Chip.

**Fig. 6.**
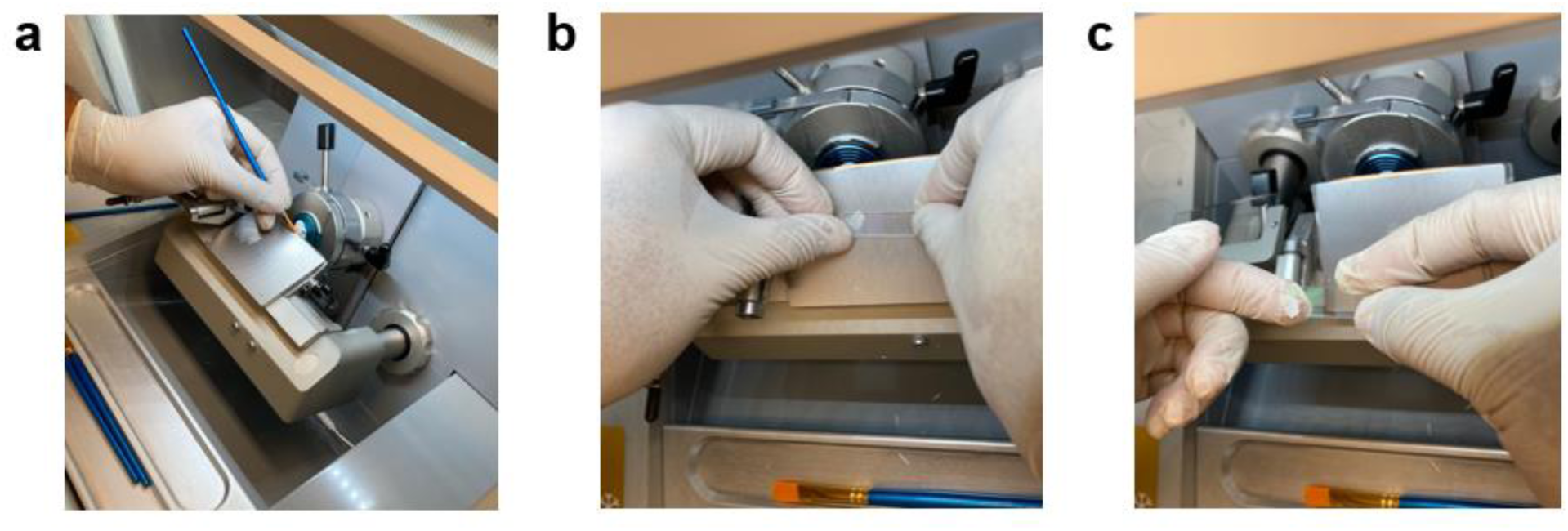
Tissue Sectioning and Attachment. (a) Cryostat set-up for sectioning. (b) Demonstration of maneuvering tissue onto the Chip surface. (c) Tissue melting for Chip attachment.

##### 2-3) [OPTIONAL] Quality check for tissue histology and RNA integrity: TIMING 1∼2 hours

1-3. Same as described in 2-2) Tissue sectioning and attachment.
4. Carefully place a glass slide over the cryostat-sectioned tissue slices, allowing the sections to adhere precisely to the slide. Preserve the slide for subsequent H&E staining and imaging to assess histological quality.
5. Slice the tissue to collect additional tissue sections (the number of sections varies according to the type of tissue and the size of the block; we typically get 3–5 sections 100 μm thick that ultimately produce adequate RNA). Return the tissue sample to the –80 °C freezer.
6. Remove the excess OCT area lacking tissues from the sections with a razor.
7. Using pre-cooled forceps, transfer the sections into a pre-cooled microcentrifuge tube.
8. Extract RNA using a Qiagen RNeasy Plus Kit according to the manufacturer’s instructions:
  a. Add 350 μL RLT buffer (supplied in the RNeasy Plus Mini Kit) with 2% beta-mercaptoethanol and 5–7 glass beads and vortex immediately for at least 30 seconds to disperse any larger tissue chunks. If large pieces are visible, cut them with scissors and/or sonicate.
  b. Centrifuge at maximum speed (15000–30000 *g*) for three minutes.
  c. Transfer the supernatant to a gDNA Eliminator spin column (supplied with the kit) placed in a collection tube. Spin for 30 seconds at 8000 *g*.
  d. Discard the column. Add 350 μL 70% EtOH to the flow-through, mix, and transfer to an RNeasy spin column placed in a collection tube. Spin for 15 seconds at 8000 *g*.
  e. Discard the flow-through. Add 700 μL RW1. Spin for 15 seconds at 8000 *g*.
  f. Discard the flow-through. Add 500 μL RPE. Spin for 15 seconds at 8000 *g*.
  g. Discard the flow-through. Add 500 μL RPE. Spin for 2 minutes at 8000 *g*.
  h. Place the column in a new collection tube and add 30–50 μL RNase-free H_2_O directly to the spin column membrane. Spin for one minute at 8000 *g* to elute the RNA.
9. Use a NanoDrop spectrophotometer to confirm that you have RNA. Subject the RNA to BioAnalyzer QC to confirm that its RIN is above 7.

##### 2-4) Tissue Fixation: TIMING 10 minutes

1. Place the Chip in a horizontal position at room temperature, surface side up (see Fig. 7a).
2. Carefully add 1 mL 4% paraformaldehyde to the surface of the Chip such that it completely covers the sections (as shown in Fig. 7b). Incubate for 10 minutes.
3. Wash the Chip with 1 mL PBS three times.

##### 2-5) Tissue H&E Staining and Imaging: TIMING ∼1 hour

1. Dispense 50 μL isopropanol into each of the four lanes/imaging units on the Chip, wait one minute, then gently tilt the Chip and dab with a paper towel to remove any residual liquid from its edges and opposite side.
2. Apply 50 μL hematoxylin across each unit, ensuring that it spreads over the tissue sections; allow it to sit for seven minutes.
3. Rinse the Chip thoroughly with 1 mL UltraPure water—covering the entire surface—to eliminate any remaining hematoxylin.
4. Lay the Chip flat and carefully add 50 μL Bluing Buffer to each unit, ensuring that it completely covers the sections within each imaging unit; leave it for two minutes.
5. Wash the Chip with 1 mL UltraPure water three times and proceed directly with the eosin staining.
6. Dispense 50 μL of Eosin Staining Solution (see below) onto each imaging area containing a tissue section. 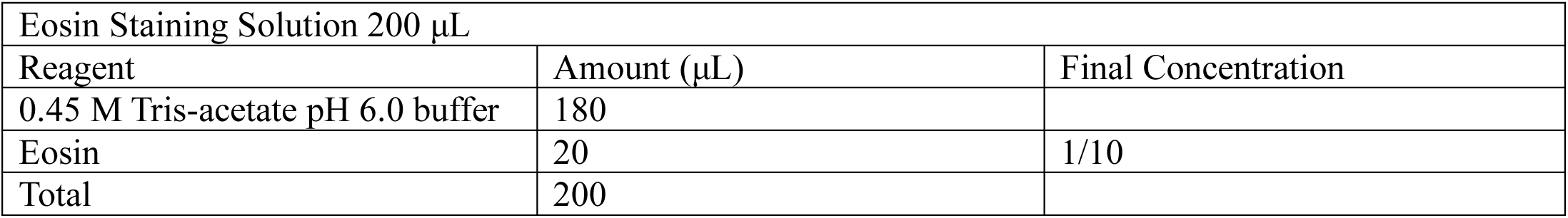
7. Aspirate the Eosin Staining Solution and wash the Chip by adding 1 mL UltraPure water to the Chip’s surface.
8. Aspirate the UltraPure water and repeat the washing step three additional times, using 1 mL UltraPure water for each rinse.
9. Aspirate all water and let the Chip dry at room temperature for 10 minutes.
10. Add 20 μL 85% glycerol to each tissue section. Carefully maneuver the Chip upside down over a coverslip measuring 25 mm × 75 mm as shown in Fig. 7c.
11. Capture images with a Keyence imager or an inverted light microscope with an upside-down position as shown in Fig. 7d. The images are later stitched together using the stitching software supplied by the Keyence imager.
12. Following the imaging step, detach the coverslip by immersing the Chip/coverslip in UltraPure water.
13. Thereafter, apply 80% ethanol to rinse away any residual glycerol, then allow it to air-dry. PAUSE POINT: The H&E-stained chip, after an 80% ethanol wash and drying, can be stored at 4°C for up to 1 week.
14. Place the Chip on a Seq-Scope adapter frame and attach the silicone isolator on the Chip as illustrated in Fig. 8a-c. The silicone isolator has adhesive on one surface that will attach tightly to the Chip surface, ensuring that enzymatic reactions can occur without contamination across the imaging area.

**Fig. 7.**
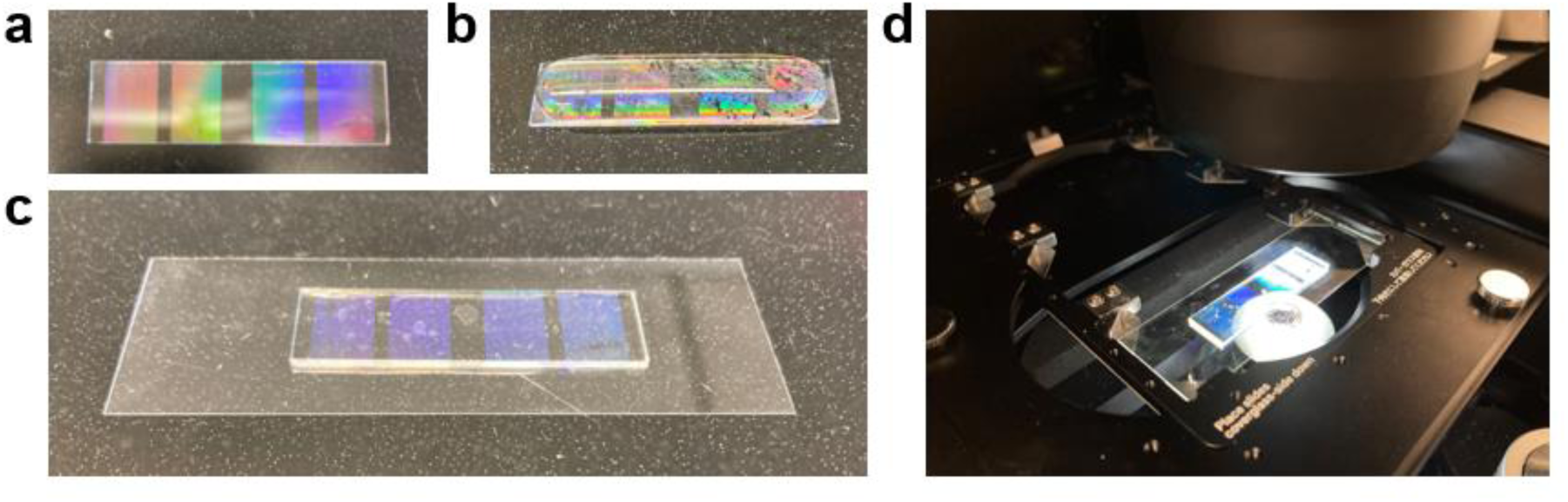
Liquid and Chip Handling Procedure in STEP 2. Tissue Fixation and Imaging Steps. The photographs are intended to serve as a guide in how to perform the procedures; therefore, actual tissues are not included in the pictures. (a) The diced Chip laid flat on the laboratory bench surface. (b) Chip covered with 1 mL of PBS. All liquid handling steps in tissue fixation and imaging will be performed similarly. The surface tension of all liquids is strong enough to hold them together on the Chip surface. (c-d) Setup of a glycerol-mounted chip/coverslip assembly on the bench (c) or in the Keyence imager (d).

**Fig. 8.**
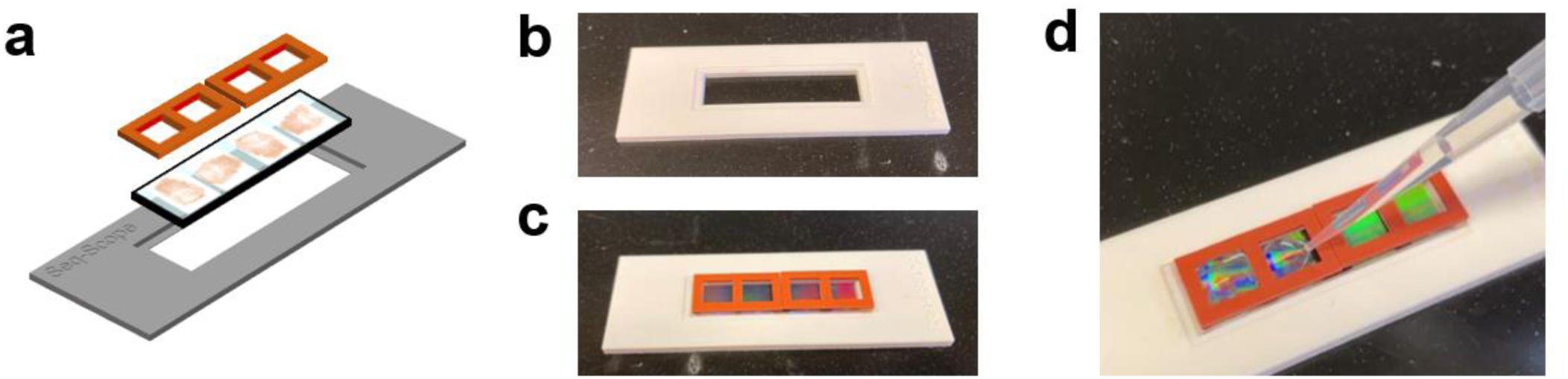
Seq-Scope Adapter Frame and Silicone Isolator used in STEP 3. Library Construction Procedures. (a) Schematic diagram showing how the Chip is placed in the Adapter Frame and how Silicone Isolators are attached to the Chip. (b) Empty Adapter Frame. (c) Adapter Frame loaded with a Chip attached to Silicone Isolators. (d) Example of how liquid handling is performed on a Seq-Scope Chip, attached by Silicone Isolators to the Adapter Frame.

#### STEP 3. Library Construction and Sequencing (2^nd^-Seq)

Steps 3-1) to 3-3) detail the procedures applicable to the entire Chip. Steps 3-4) to 3-7) describe scalable procedures that apply to each imaging area of the Chip, allowing them to be conducted independently or simultaneously, depending on the user’s preference.

##### 3-1) mRNA Release and Reverse Transcription: TIMING 16∼20 hours

1. Add 1 μL Collagenase I stock solution into pre-warmed HBSS to make the Collagenase Working Solution. Thoroughly mix using a pipette and pre-warm to 37 °C. 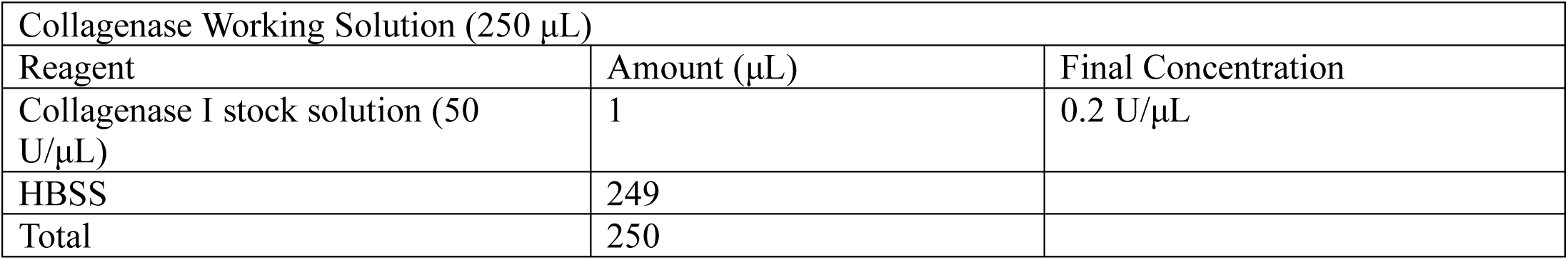
2. Dispense 50 μL Collagenase Working Solution onto each unit of the Chip
3. Incubate at 37 °C for 20 minutes.
4. Add 2 μL Pepsin stock solution into 0.1M HCl to make the Pepsin Working Solution. Thoroughly mix using a pipette and pre-warm to 37 °C. 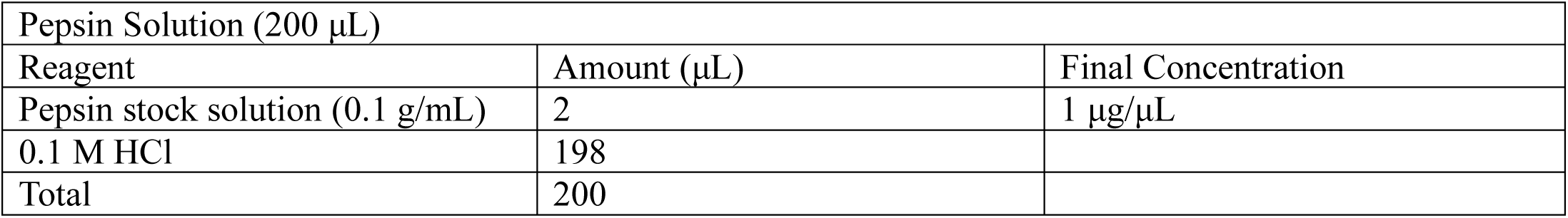
5. Dispense 50 μL Pepsin Working Solution onto each imaging unit of the Chip and incubate at 37 °C for 10 minutes. CRITICAL: Optimization of pepsin-treatment time is critical for transcript capture and library construction efficiency in the subsequent procedures. Ten minutes works well for mouse liver, but different tissue types will warrant different digestion times. To optimize these conditions, you can use either the method described in the original spatial transcriptomics paper^15^ or the tissue optimization kit commercially available for the 10x Genomics Visium platform^16^. We found that the conditions optimized for low-resolution ST methods also works well for high-resolution Seq-Scope. Moreover, it is feasible to test the optimization of the permeabilization conditions directly within the Seq-Scope workflow, given its cost-effectiveness for low-depth sequencing. CRITICAL: If there are concerns about the accuracy of pipetting, particularly when dispensing multiple 50 μL aliquots from a 200 μL total solution, it is advisable to adjust the dispensed volume down to 48 μL. This optional adjustment helps ensure that the final aliquot is not substantially less than intended, compensating for potential pipetting discrepancies.
6. Set up the RT Equilibration Solution using the following formulation. 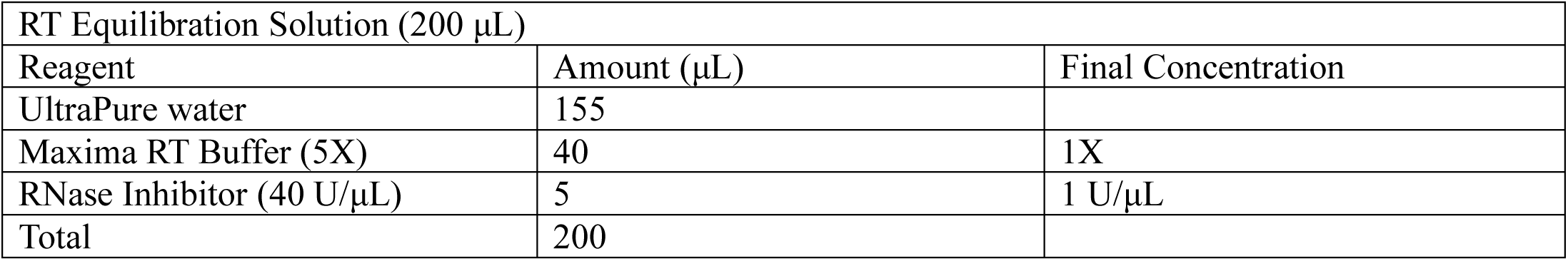
7. Dispense 50 μL RT Equilibration Solution onto each unit of the Chip. Incubate at room temperature for 10 minutes.
8. Set up the RT Solution using the following formulation. 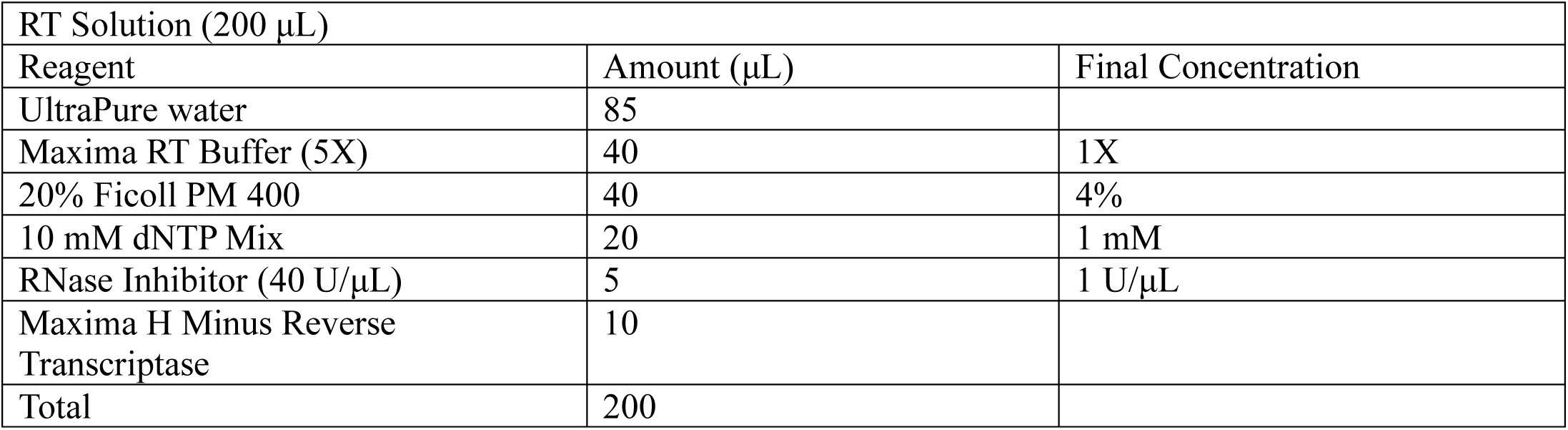
9. Dispense 50 μL RT Solution onto each section of the Chip. Prepare a humidified chamber as detailed in Steps 1–3. In the present setup, place the Seq-Scope Chip—now loaded with RT solution—above the Parafilm in an upright position. Seal the chamber securely and allow the setup to incubate overnight at 42 °C. CRITICAL: Verify that the humidified chamber is properly sealed and ensure that the RT Solution remains moist, avoiding evaporation during the overnight incubation.
10. After the overnight incubation, remove the RT solution.

##### 3-2) Tissue Removal: TIMING 1 hour 30 minutes

1. Prepare an additional 200 μL of Exonuclease I Solution as described in Step 1-3).
2. Dispense 50 μL Exonuclease I Solution onto each unit of the Chip. Incubate for 45 minutes at 37 °C in a humidified chamber. This step eliminates all capture probes that failed to capture mRNA.
3. Set up the Tissue Digestion Buffer using the following formulation. 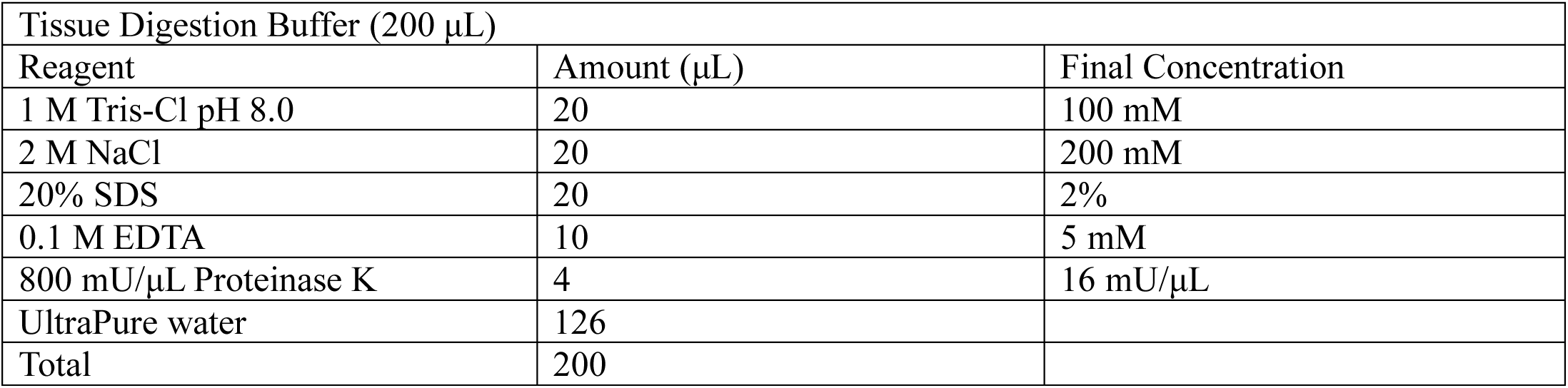
4. Dispense 50 μL Tissue Digestion Buffer onto each unit of the Chip. Incubate for 45 minutes at 37 ℃ in a humidified chamber. This step removes all tissue remnants except mRNAs bound to capture molecules.
5. Rinse each unit of the Chip sequentially with the following solutions.
  a. 200 μL UltraPure water (3 times, incubate 1 minute each at room temperature)
  b. 200 μL 0.1N NaOH (3 times, incubate 5 minute each at room temperature)
  c. 200 μL 0.1M Tris pH 7.5 (3 times, incubate 1 minute each at room temperature)
  d. 200 μL UltraPure water (3 times, incubate 1 minute each at room temperature)
6. The Chip now contains HDMI-tagged cDNA molecules on its surface and is ready for secondary strand synthesis.

#### 3-3) Secondary strand synthesis: TIMING 2 hour 30 minutes

1. Set up the Random Priming and Extension Mix (RPEM) using the following formulation. 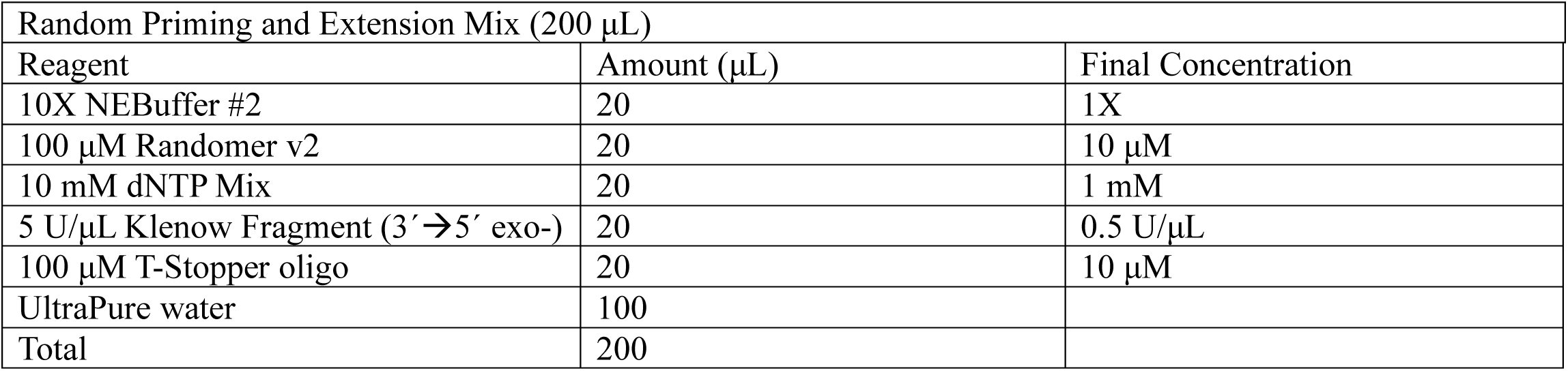 CRITICAL: Compared to Randomer v1 described in our original paper^9^, Randomer v2 contains a non-A nucleotide at the terminal position, suppressing the synthesis of poly-A-only secondary strands that would generate data without any useful information such as 3’-UTR or coding sequences. CRITICAL: The addition of T-Stopper is optional; however, we found that it further suppresses the formation of poly-A-only secondary strands by scavenging Randomers preferring oligo-dT sequences during secondary strand synthesis.
2. Dispense 50 μL RPEM onto each unit of the Chip. Incubate for two hours at 37 ℃ in a humidified chamber.
3. After incubation, rinse each unit of the Chip using 200 μL UltraPure water. Repeat three times.
4. Dispense 50 μL 0.1N NaOH onto each unit of the Chip. Incubate for five minutes at room temperature.
5. Collect the 50 μL NaOH solution containing the secondary strand and transfer it to a 1.5 mL LoBind tube.
6. Repeat the secondary strand extraction procedure (4–5) to collect another 50 μL of NaOH solution containing secondary strand products from each unit (a total of 100 μL for each unit).
7. Add 50 μL neutralization buffer (Zymogen, D4027-3-50) to each 100 μL of NaOH solution containing the secondary strand products. Mix well using a pipette.
8. Adjust the volume of the crude secondary strand solution to precisely 150 μL using UltraPure water.
9. Proceed directly to the purification step without delay. All subsequent procedures are specified for individual imaging units.

#### 3-4) Purification of the secondary strand solution: TIMING ∼30 minutes

1. Prepare 10 mL of fresh 80% (vol/vol) ethanol in a 15 mL conical tube by mixing 8 mL 100% ethanol and 2 mL UltraPure water. Thoroughly mix the solution by vortexing vigorously for 10 seconds.
2. Pre-warm AMPure XP magnetic beads to room temperature. Completely resuspend the beads by vortexing vigorously for one minute.
3. Transfer 180 μL of fully resuspended beads to the tube containing 150 μL of crude secondary strand solution. Mix well by pipetting up and down more than 10 times.
4. Let the mixture sit at room temperature for 10 minutes to allow for effective interaction between the magnetic beads and the solution.
5. Position the tube containing the suspension on a magnetic stand that accommodates 1.5-mL tubes and leave it for five minutes to allow the magnetic beads to separate from the solution effectively.
6. Carefully aspirate and discard the supernatant, taking care to leave the magnetic beads undisturbed at the bottom of the tube.
7. While maintaining the tube on the magnetic stand, gently wash the beads by adding 1 mL of freshly prepared 80% ethanol to the tube, using caution not to disturb the magnetic bead pellet.
8. After adding the ethanol, allow the tube to incubate on the magnetic stand for 30 seconds, then carefully remove and discard the ethanol supernatant, ensuring that the bead pellet remains undisturbed.
9. Perform a second wash by adding 1 mL of freshly prepared 80% ethanol to the beads, following the same procedure as the previous wash to ensure thorough cleaning.
10. With the tube still on the magnetic stand, carefully aspirate and dispose of any remaining ethanol supernatant, ensuring that the magnetic beads remain undisturbed at the bottom of the tube.
11. With the tube remaining on the magnetic stand, allow the magnetic beads to air-dry at room temperature until the residual ethanol has completely evaporated and the beads appear dry. Avoid over-drying as this can negatively affect bead functionality. Typically, air-drying requires approximately five minutes; it is important to visually inspect the beads for a matte appearance rather than glossy to ensure adequate drying.
12. Once the beads have air-dried to a matte appearance, remove the tube from the magnetic stand and add 80 μL UltraPure water. Resuspend the beads by pipetting up and down at least 10 times until they are fully suspended.
13. Allow the sample to sit at room temperature for two minutes. Should any droplets adhere to the sides of the tube, centrifuge briefly to consolidate the liquid at the bottom of the tube.
14. Rest the tube on the magnetic stand until the solution becomes clear—typically ∼30 seconds.
15. Carefully transfer the clear supernatant to new tubes using a pipette.
16. This supernatant constitutes the purified secondary strand solution.

PAUSE POINT: The purified secondary strand can be stored at –20°C for up to 6 months.

#### 3-5) Library Amplification PCR reaction and Purification: TIMING ∼3 hours

1. Prepare the Library Amplification PCR mix according to the formulation provided below. 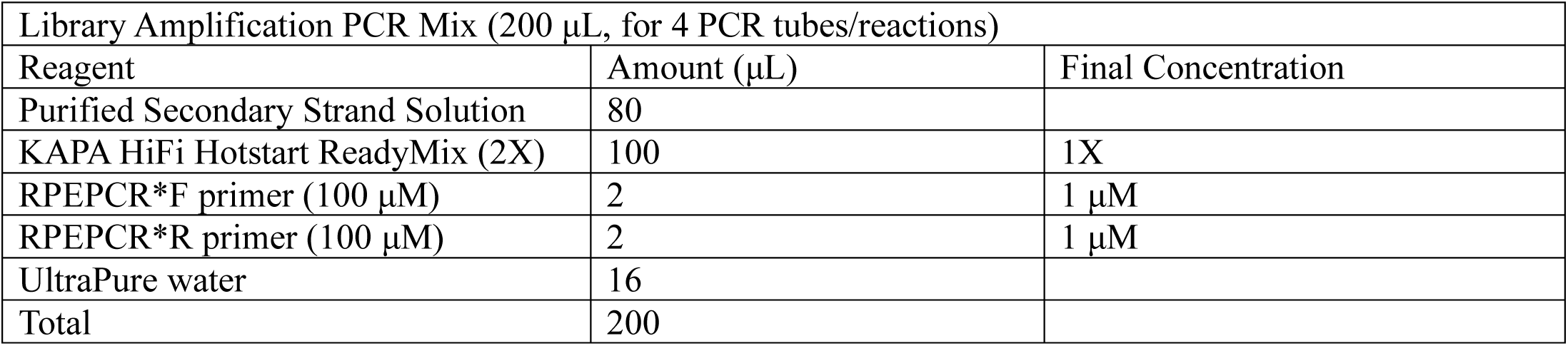
2. Distribute the Library Amplification PCR mix evenly into four PCR tubes, with 50 μL of the mix in each tube. Proceed to PCR using the conditions described below:
  a) Initial Denaturation: 3 minutes at 95°C, performed once.
  b) Cycling Steps (repeat 13–15 times):
    - Denaturation: 30 seconds at 95 °C.
    - Annealing: 1 minute at 60 °C.
    - Extension: 1 minute at 72 °C.
  c) Final Extension: 2 minutes at 72 °C, performed once.
  d) Hold: Indefinitely at 4 °C.
3. Following PCR completion, briefly centrifuge the PCR tubes to collect any condensation. Then pool the contents into one 1.5 mL LoBind tube and adjust the total volume to 200 μL with UltraPure water if necessary.
4. Pre-warm AMPure XP magnetic beads to room temperature. Completely resuspend the beads by vortexing vigorously for one minute.
5. Transfer 200 μL of the fully resuspended beads to the tube containing 200 μL of crude secondary strand solution. Mix well by pipetting up and down more than 10 times.
6. Let the mixture sit at room temperature for five minutes to allow effective interaction between the magnetic beads and the solution.
7. Position the tube containing the suspension on a magnetic stand that accommodates 1.5-mL tubes and leave it for three minutes to enable the magnetic beads to separate from the solution effectively.
8. Carefully aspirate and discard the supernatant, using caution to leave the magnetic beads undisturbed at the bottom of the tube.
9. While maintaining the tube on the magnetic stand, gently wash the beads by adding 200 μL of freshly prepared 80% ethanol to the tube, being careful not to disturb the magnetic bead pellet.
10. After adding the ethanol, allow the tube to incubate on the magnetic stand for 30 seconds, then carefully remove and discard the ethanol supernatant, ensuring that the bead pellet remains undisturbed.
11. Perform a second wash by adding 200 μL of freshly prepared 80% ethanol to the beads, following the same procedure as the previous wash to ensure thorough cleaning.
12. With the tube still on the magnetic stand, carefully aspirate and dispose of any remaining ethanol supernatant, ensuring that the magnetic beads remain undisturbed at the bottom of the tube.
13. With the tube remaining on the magnetic stand, allow the magnetic beads to air-dry at room temperature until the residual ethanol has completely evaporated and the beads appear dry. Avoid over-drying as this can affect bead functionality. Typically, air-drying requires approximately five minutes; it is important to visually inspect the beads for a matte appearance rather than glossy to ensure adequate drying.
14. Once the beads have air-dried to a matte appearance, remove the tube from the magnetic stand and add 40 μL of UltraPure water. Resuspend the beads completely by pipetting up and down at least 10 times.
15. Allow the sample to sit at room temperature for two minutes. Should any droplets adhere to the sides of the tube, centrifuge briefly to consolidate the liquid at the bottom.
16. Rest the tube on the magnetic stand until the solution becomes clear—typically ∼30 seconds.
17. Carefully transfer the clear supernatant (40 μL) to fresh tubes using a pipette.
18. This supernatant constitutes the purified Library Amplification PCR product, ready for subsequent Indexing PCR.
19. Measure the eluate’s concentration with a Qubit fluorometer.
20. [OPTIONAL] Evaluate the Library Amplification PCR product by either agarose gel electrophoresis or BioAnalyzer. Generally, these methods should generate a broad smear ranging from approximately 200 bp to 2000 bp, indicating a diverse range of fragment sizes. The observed trace should be a continuous smear—devoid of any distinct, ladder-like bands—indicating a uniformly amplified library with a broad range of PCR products.
21. Calculate the library’s molar concentration using the formula: (*concentration in ng/μl*) × 3 = *concentration in nM* This simplified formula is based on the assumption that the average library size is 500 bp and that the average molecular weight of a DNA base pair is 660 g/mol. The outcome, expressed in nM, is crucial for determining the volume of template needed for the subsequent indexing PCR step.
22. Store the prepared Library Amplification PCR products in a –20 °C freezer. CRITICAL STEP: To ensure the success of the experiments, the library concentrations should ideally fall between 10 and 30 ng/μL. Nonetheless, libraries with lower concentrations can still proceed, provided they are capable of being amplified in the indexing PCR. PAUSE POINT: The purified Library Amplification PCR product can be stored at –20°C for up to 6 months.

#### 3-6) Indexing PCR and Size Selection: TIMING ∼2 hours

1. If higher than 2 nM, dilute the purified Library Amplification PCR product to 2 nM.
2. Set up the Indexing PCR mix using the following formulation. 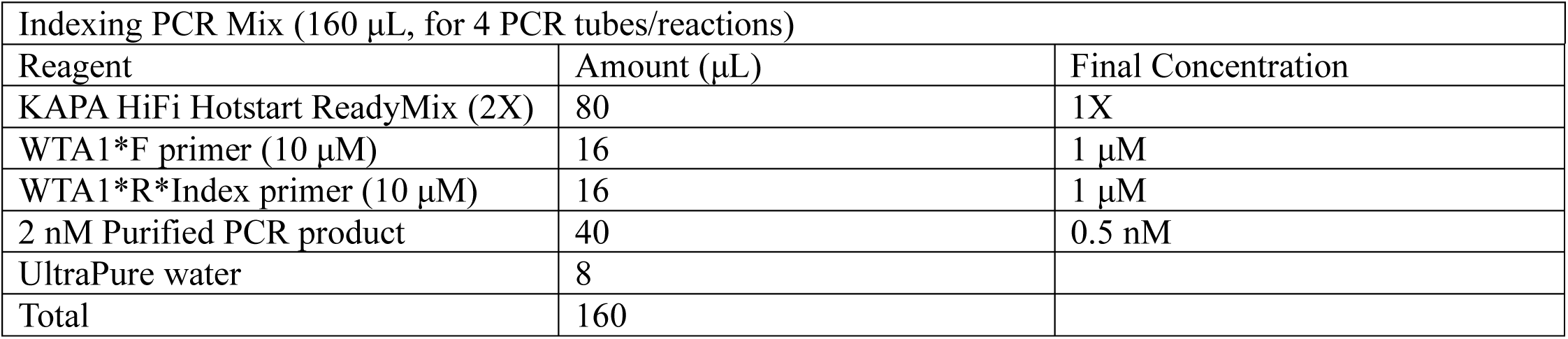
3. Perform Indexing PCR in 4 PCR tubes, each with 40 μL volume, using the following conditions:
  a) Initial Denaturation: 3 minutes at 95 °C, performed once.
  b) Cycling Steps (repeat 8–9 times):
    - Denaturation: 30 seconds at 95 °C.
    - Annealing: 30 seconds at 60 °C.
    - Extension: 30 seconds at 72 °C.
  c) Final Extension: 2 minutes at 72 °C, performed once.
  d) Hold: Indefinitely at 4 °C.
4. Combine the 4 indexing PCR reactions into a new 1.5-mL LoBind tube.
5. Using the Zymogen DNA Clean and Concentration-25 kit, purify and concentrate the combined indexing PCR products—following the manufacturer’s protocol—and elute in 40 μL of UltraPure water.
6. Add 8 μL of 6X Gel Loading Dye Purple to the 40 μL of concentrated indexing PCR product. Mix thoroughly using a pipette.
7. Load the prepared sample onto a 2% agarose gel and proceed with electrophoresis, continuing until the purple dye has migrated to the bottom of the gel. CRITICAL: Be sure to clean the agarose gel tank and imaging station thoroughly to avoid contamination. Always use a fresh blade for slicing the gel.
8. Stain the gel by soaking it in 1X SYBR™ Gold Nucleic Acid Gel Staining solution (made by diluting the 10,000X stock with the gel running buffer). Incubation time varies depending on the gel volume.
9. Using a blue light illuminator, excise the gel segment containing DNA fragments sized between 400 and 850 bp.
10. Perform gel extraction with the Zymogen Gel DNA Recovery kit according to the manufacturer’s protocol. In the final step, elute the DNA in 100 μL of UltraPure water in a LoBind 1.5-mL tube.
11. Pre-warm AMPure XP magnetic beads to room temperature. Completely resuspend the beads by vortexing vigorously for one minute.
12. Transfer 70 μL of the fully resuspended beads to the tube containing 100 μL of crude secondary strand solution. Mix well by pipetting up and down more than 10 times.
13. Let the mixture sit at room temperature for five minutes to allow effective interaction between the magnetic beads and the solution.
14. Position the tube with the suspension on a magnetic stand that accommodates 1.5-mL tubes and leave it for three minutes to enable the magnetic beads to separate from the solution effectively.
15. Carefully aspirate and discard the supernatant, taking care to leave the magnetic beads undisturbed at the bottom of the tube.
16. While maintaining the tube on the magnetic stand, gently wash the beads by adding 200 μL of freshly prepared 80% ethanol to the tube, being sure not to disturb the magnetic bead pellet.
17. After adding the ethanol, allow the tube to incubate on the magnetic stand for 30 seconds, then carefully remove and discard the ethanol supernatant, ensuring the bead pellet remains undisturbed.
18. Perform a second wash by adding 200 μL of freshly prepared 80% ethanol to the beads, following the same procedure as the previous wash, to ensure thorough cleaning.
19. With the tube still on the magnetic stand, carefully aspirate and dispose of any remaining ethanol supernatant, ensuring the magnetic beads remain undisturbed at the bottom of the tube.
20. With the tube remaining on the magnetic stand, allow the magnetic beads to air-dry at room temperature until the residual ethanol has completely evaporated and the beads appear dry. Avoid over-drying as this can affect bead functionality. Typically, air-drying requires approximately five minutes; it is important to visually inspect the beads for a matte appearance rather than glossy to ensure adequate drying.
21. Once the beads have air-dried to a matte appearance, remove the tube from the magnetic stand and add 20 μL of UltraPure water. Resuspend the beads completely by pipetting up and down at least 10 times.
22. Allow the sample to sit at room temperature for 2 minutes. Should any droplets adhere to the sides of the tube, centrifuge briefly to consolidate the liquid at the bottom.
23. Rest the tube on the magnetic stand until the solution becomes clear—typically ∼30 seconds.
24. Carefully transfer the clear supernatant (20 μL) to fresh tubes using a pipette.
25. Determine the concentration of the eluate using a Qubit fluorometer.
26. Store the final library preparations at –20 °C; they remain stable and suitable for sequencing for up to six months.

PAUSE POINT: The final library can be stored at –20°C for up to 6 months.

#### 3-7) Library Sequencing

1. Quantitate the library via BioAnalyzer or LabChip—i.e., a fragment analysis platform that performs well on longer libraries (TapeStation is not advised). The band should appear between 400 and 850 bp, consistent with the size selection scheme. CRITICAL: If contaminant bands are identified at molecular weights below 300 bp, gel running-based size selection or 0.7x AMPure XP bead purification should be further repeated to clean the library. It is critical to remove adapter dimers and abnormal products to ensure high-quality sequencing.
2. [OPTIONAL] A SequencingQC method involving MiSeq Nano flow cells can be implemented to predict the clustering behavior of the library. The read-count data from these diagnostic runs reveal whether the library is likely to cluster well on larger NovaSeq flow cells. CRITICAL: Likely due to the presence of extended conserved sequences in the library, qPCR using kits designed to quantitate Illumina-adapter-ligated products returned results showing reduced library concentration, leading to underestimation and overloading of the library. Therefore, appropriate loading concentration is determined more effectively via Qubit with BioAnalyzer/LabChip quantification alone or in conjunction with the SequencingQC method.
3. The library can be sequenced using most Illumina sequencing configurations, including PE150 or PE100. However, only 32 bp of Read 1 are used for spatial barcoding. Custom configurations are also possible; in such cases, at least 32 cycles for Read 1, 50 cycles for Read 2, and 8 cycles for Index 1 (if multiplexing) must be sequenced. For example, sequencing can be performed using a NovaSeq-X 1.5B 100 cycle kit (with 138 available cycles), allocating 32 cycles to Read 1, 8 cycles to Index 1, and 98 cycles to Read 2. Initial testing can begin with a shallow read depth (e.g., 100 million reads); if the spatial capture pattern is satisfactory, deeper sequencing can be performed to uncover more comprehensive information.

## B. Computational Procedures

The analysis of sequence reads is divided into three key steps (Fig. 9): Step 1 builds the map of spatial barcodes (HDMI) from the single-ended sequence reads of the chip (1^st^-Seq). Step 2 generates a high-resolution spatial expression (SGE) matrix, by aligning the paired-end sequence reads of the prepared library (2^nd^-Seq) by aligning the cDNA sequence while resolving the spatial positions of the HDMI barcodes. Optionally, if histology images are available, they can also be aligned to the HDMI coordinates in this step. Step 3 describes an exemplary downstream analysis of the SGE matrix, including the identification of cell type factors and the inference of pixel-level factors at submicrometer resolution. Steps 1 and 2 are implemented in the NovaScope pipeline (https://github.com/seqscope/NovaScope), which is an extension of the STtools pipeline^19^. Step 3 primarily uses the FICTURE software tool^29^ and the detailed steps to reproduce our exemplary analysis is provided in a public GitHub repository (https://github.com/seqscope/NovaScope-exemplary-downstream-analysis). Our aligned sequenced reads can be directly used for tasks that require read-level information, such as allele-specific expression or somatic variant analysis. Our SGE matrix can also be analyzed with many additional tools, such as Giotto and Squidipy, offering unlimited possibilities for downstream data analysis.

**Fig. 9.**
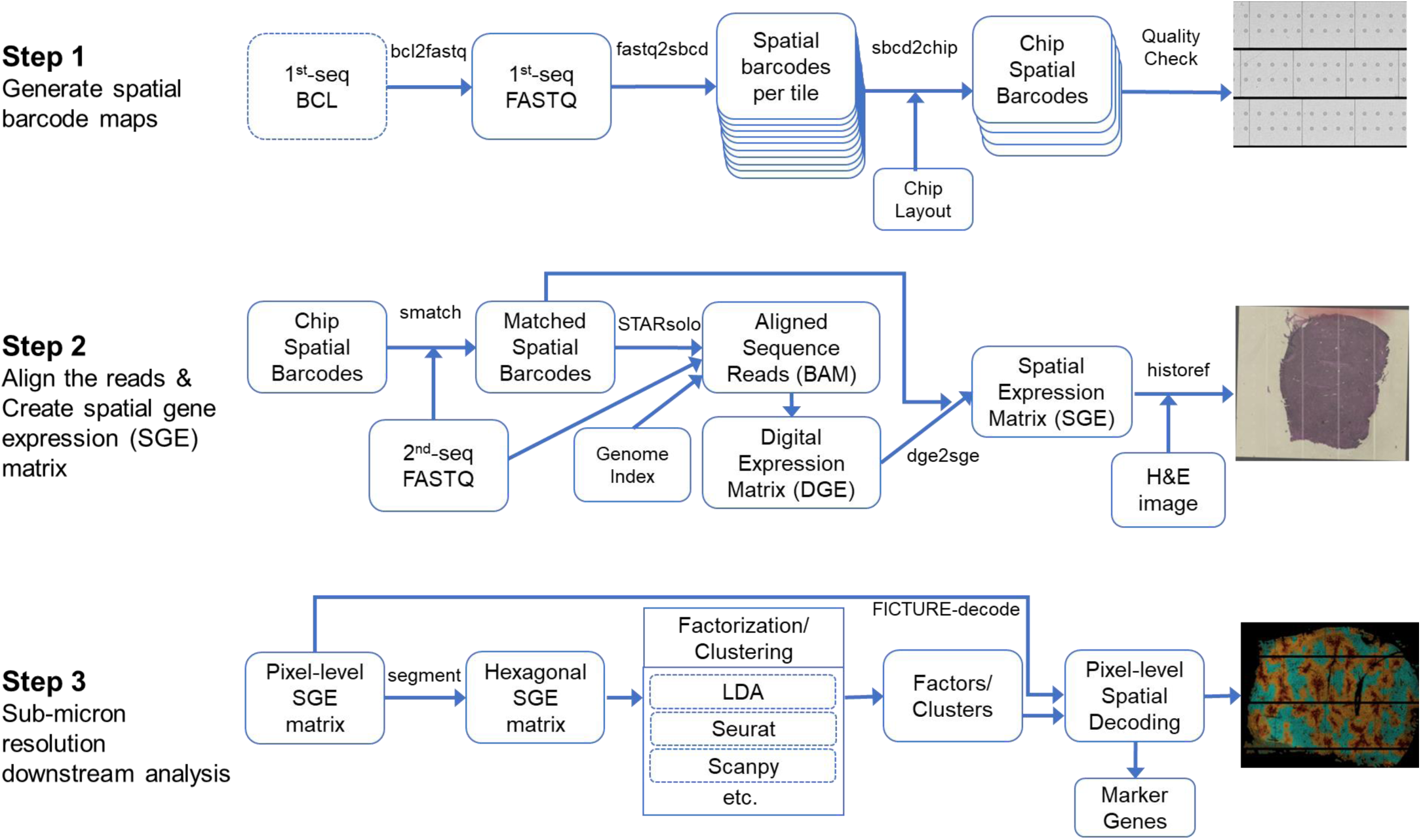
Overview of Computational Procedures.

### STEP 1. Generation of Spatial Barcode Maps

#### 1-1) Install NovaScope pipeline

CRITICAL: It is important to ensure that all dependent software tools are installed before installing and testing the NovaScope pipeline.

1. Navigate to https://github.com/seqscope/novascope
2. Following the instruction, clone the GitHub repository
3. If needed, install dependent software tools, such as Snakemake and ImageMagick, as indicated in the documentation.
4. Validate the installation. Test the pipeline with the example dataset provided in the repository.

#### 1-2) (Optional) Generate 1st-seq FASTQs from BCL files

1. If FASTQ files of 1^st^-Seq were delivered from the sequencing center, this procedure can be skipped.
2. If only BCL files are available, convert the corresponding Illumina BCL-formatted files into FASTQ files using the *bcl2fastq* tool. In the output directory, a single-ended FASTQ file for each of the four lanes will be generated.

#### 1-3) Build a spatial barcode map for individual tiles

Please note that steps 1-3) to 2-3) can be performed all at once, using “all” command in NovaScope, which will make the procedure more streamlined. This option is described in step 2-4).

1. Using the *fastq2sbcd* step in NovaScope, generate a map of spatial barcodes for each tile. The spatial barcodes (HDMI, typically 32 bp) and their spatial coordinates (lane, tile, X, Y positions) are extracted from the FASTQ file, reverse-complemented for easier match to the 2nd-Seq sequences and stored by individual tiles. The output of this step includes compressed tab-delimited files containing a mapping between barcodes and spatial coordinates for each tile and a summary file across all tiles.
2. Check the summary statistics from the **`manifest.tsv`** file from the output directory. Each lane should contain 936 tiles, and each tile typically contains 3 million or more reads. Most of the reads should match to the expected patterns of HDMI, and the **`manifest.tsv`** file reports how many reads matches to the expected pattern.
3. Check the statistics of unique and duplicate barcodes to assess the quality of sequence reads using the summary output file. A lane with 80% or less unique reads (i.e., 20% or more duplicate reads) typically suggest underclustering, which would require additional titration of library loading concentration (see TROUBLESHOOTING).

#### 1-4) Build a spatial barcode map for each Chip

1. Using the *sbcd2chip* step in NovaScope, merge the spatial barcodes for a section chip across tiles based on the layout of those tiles in the flow cell. A total of 936 tiles from a single lane are arranged in six swaths, with each swath containing 78 tiles, displayed on both the top and bottom surfaces. Their spatial coordinates are established by aligning a histology image with spatial barcodes, using the fiducial marks as anchors (Fig. 10). Utilizing the specified gaps and shifts between tiles, as provided with NovaScope, a map of spatial barcodes will be generated for each chip section, following the removal of duplicated barcodes within the chip area. The output of this step comprises a compressed, tab-delimited file that contains a mapping between barcodes and global spatial coordinates within the chip. This mapping merges data across all tiles, following the removal of duplicate barcodes. Additionally, the output includes an image that displays the distribution of the spatial coordinates of the barcodes.
2. Examine the image of the merged spatial barcode map of the Chip (Fig. 11). Because the fiducial marks are deleted for sequence reads, they should look empty, and they should be aligned with other fiducial marks horizontally and vertically.
3. Check the number of unique barcodes from the summary output file.

**Fig. 10.**
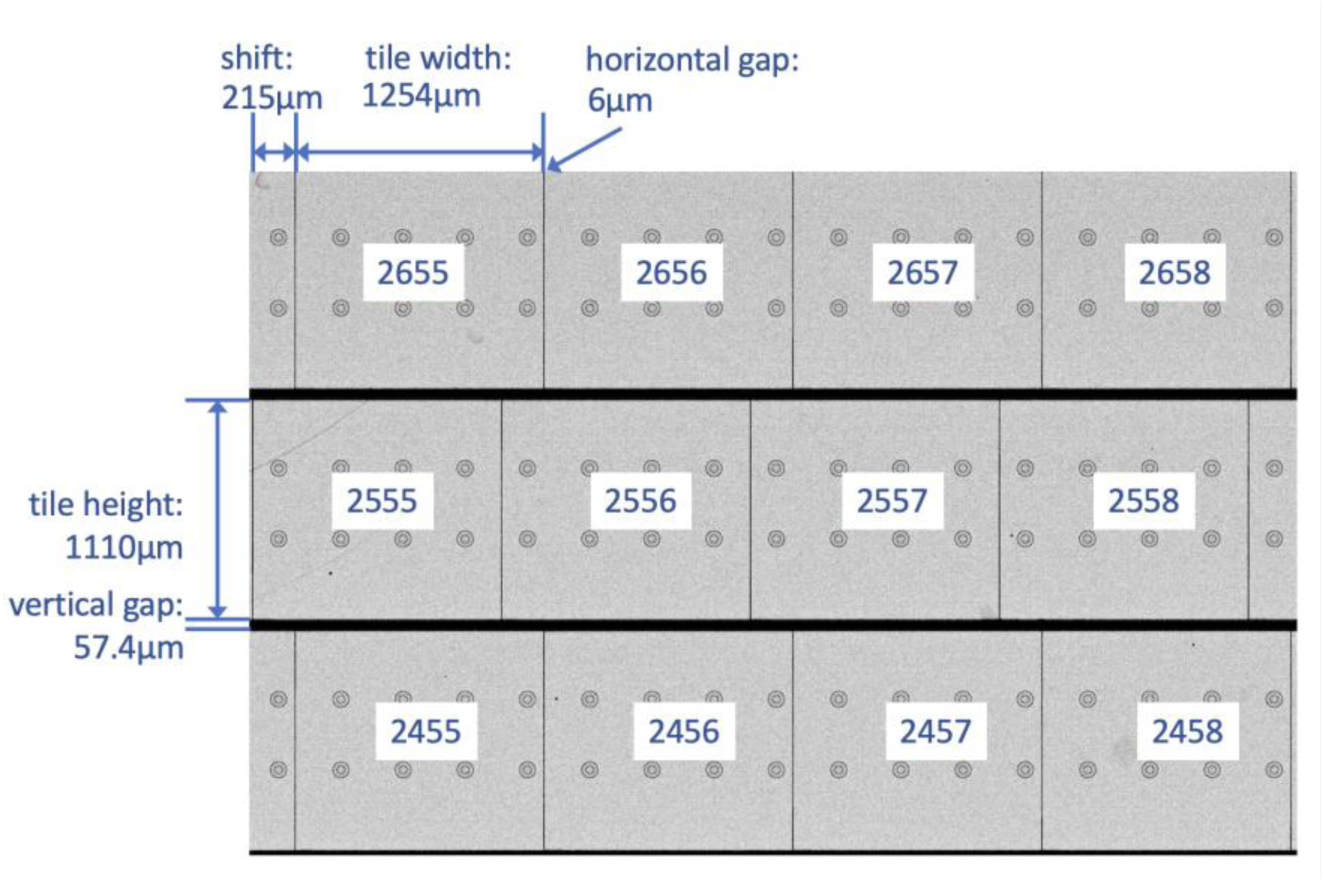
Spatial arrangement between tiles, which is determined by comparing the histology images and the spatial barcodes.

**Fig. 11.**
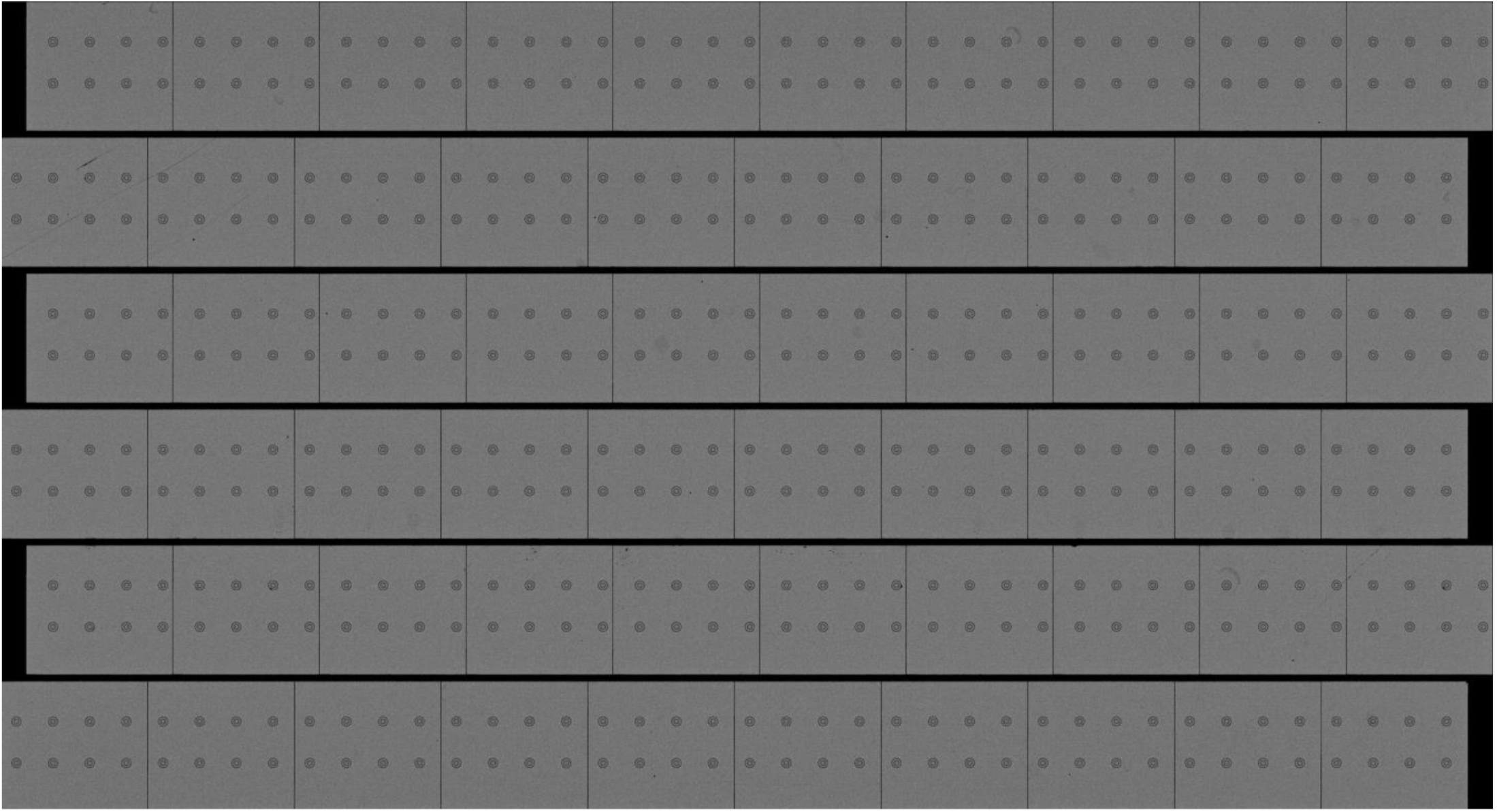
An output sbcd image from step 1-4), showing the distribution of spatial barcodes (i.e., HDMIs). The fiducial marks should be aligned horizontally and vertically with correct parameter settings.

### STEP 2. Sequence Alignment and Spatial Gene Expression (SGE) Matrix Generation

#### 2-1) Match 2^nd^-Seq reads to spatial barcodes of the corresponding Chip

1. For each FASTQ file containing 2^nd^-seq reads, determine whether the HDMI sequences in read 1 are found from the spatial barcodes in the Chip, using the *smatch* step of NovaScope. The output of this step includes a compressed tab-delimited file containing spatial barcodes matching the 2^nd^-seq reads, an image of the distribution of the spatial coordinates of the matching barcodes, and a file summarizing the count of 2^nd^-seq reads based on the matching results.
2. Check the output summary metrics to ensure that the matching rate is not too low (e.g., < 5%). If the matching rate is too low, double-check whether the sample swap has happened.
3. Check the output image that visualizes the spatial coordinates of matching barcodes, which should roughly match the tissue area (Fig. 12). If the matching barcodes do not show an expected pattern, check if there were any issues in experimental procedures, such as unsuccessful tissue permeabilization. See TROUBLESHOOTING for details.

**Fig. 12.**
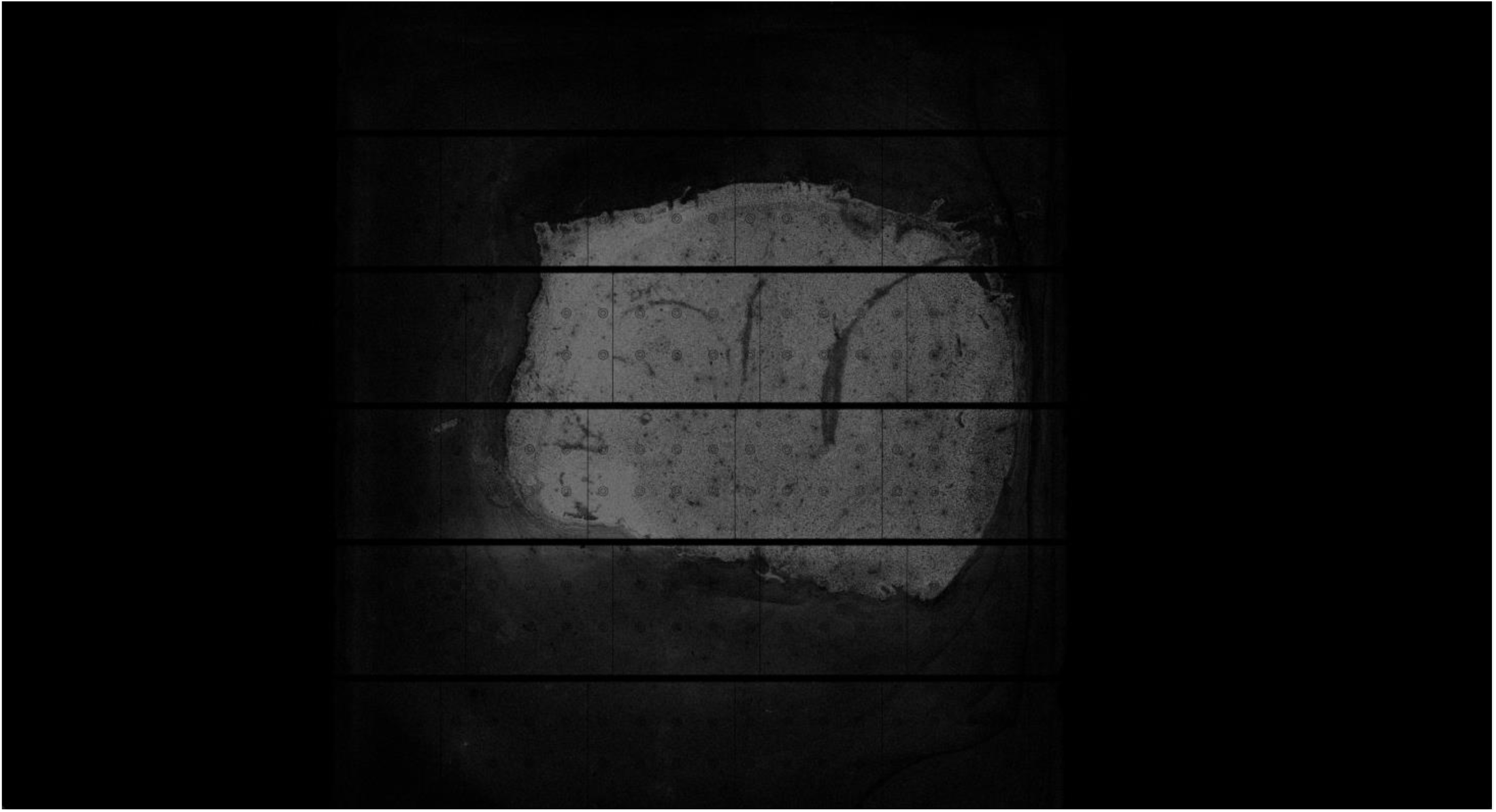
An output smatch image from Step 2-1), showing the distribution of spatial barcodes (i.e., HDMIs) that matched to the 2^nd^-seq FASTQ files. The distribution should be consistent to the tissue area.

#### 2-2) Align the 2^nd^-Seq reads to the reference genome and generate an SGE matrix

1. Using the *align* step in NovaScope, combine all 2^nd^-Seq FASTQ files sequenced for the Chip, filter out reads without HDMI sequence matching the Chip’s spatial barcodes, and align the reads to the reference genome with STARsolo. Then, utilize the *dge2sdge* step in NovaScope to generate a spatial gene expression (SGE) matrix by joining the STARsolo output and spatial barcode dictionary. The output of this step includes a BAM file of aligned reads, sorted by genomic coordinates, generated from STARsolo. It also includes the SGE matrix, compatible with the MatrixMarket format used in 10x Genomics output. Additional images and summary metrics are also generated.
2. Check the output files from the pipeline to ensure that the total number of reads, proportion of reads aligned to genomes and genes, library saturation, the number of aligned spatial barcodes, and the number of unique transcripts.
3. Check the output image that visualizes the spatial distribution of aligned transcripts to see if it matches to the tissue area and if it is consistent with the spatial distribution of matching barcodes (Fig. 13).

**Fig. 13.**
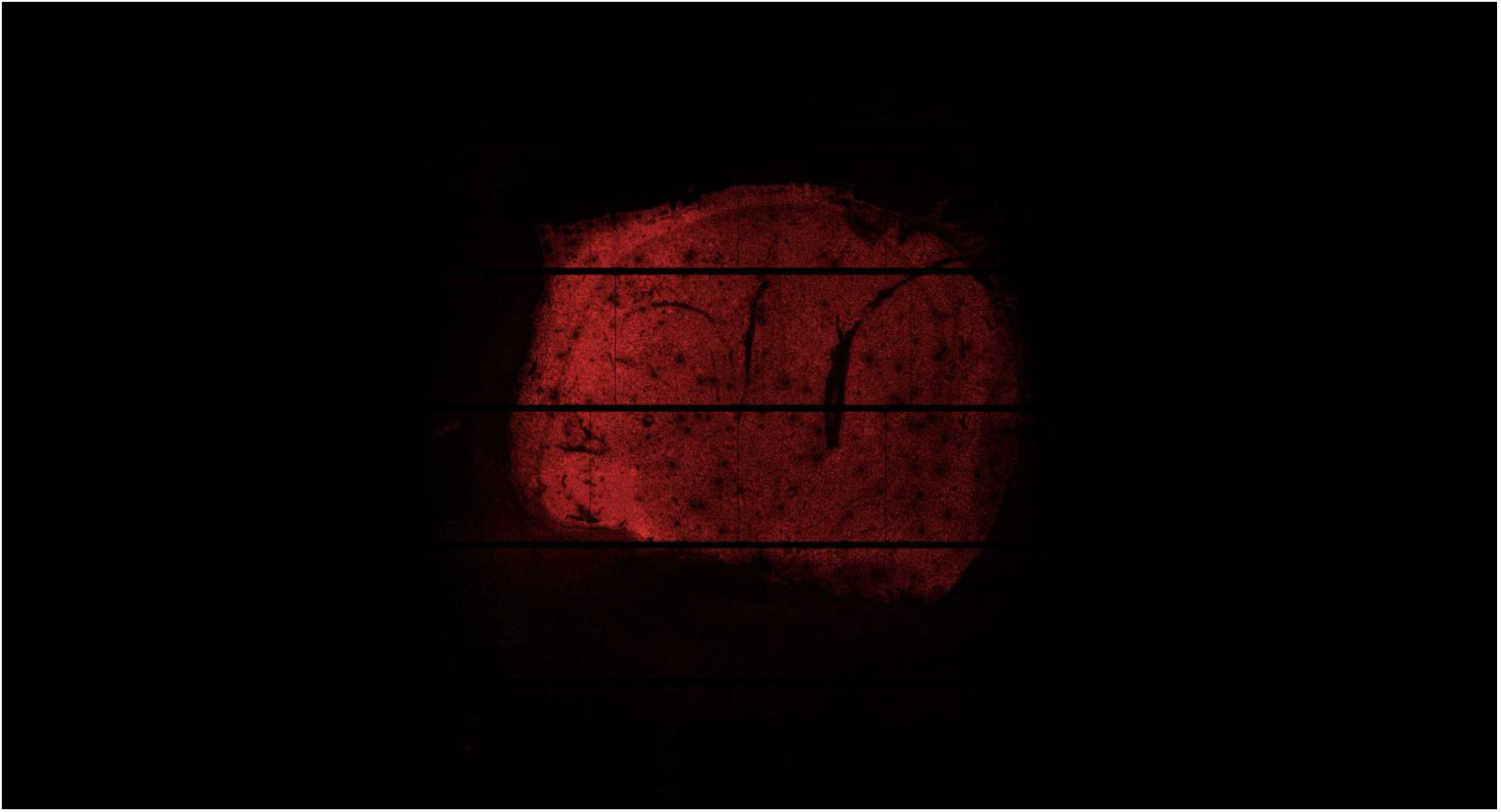
An output sge image from step 2-2), showing the distribution of spatial barcodes (i.e., HDMIs) that aligns to the reference genome, coloring exon-aligned transcripts as red, unspliced transcripts as green, and mitochondrial transcripts as blue.

#### 2-3) (Optional) Histology alignment based on fiducial mark and tissue density

1. If an H&E image is available, use the *historef* step in NovaScope to align the image to the spatial coordinates of the SGE matrix. The alignment is performed by matching fiducial marks that are visible from both images, minimizing the difference between them. The output of this step includes a referenced geotiff file that allows the coordinate transformation between the SGE matrix and histology image and another tiff file that has the exact same dimension as the output image from steps 2-1) and 2-2), in addition to several QC reports.
2. Check the output files to ensure that smatch (Fig. 12), sge (Fig. 13), and histology (Fig. 14) images match precisely. If the fiducial marks look clear from both images, the match should be accurate in submicrometer resolution when overlaid. However, in cases where an insufficient number of fiducial marks are visible and/or the fiducial marks were aligned incorrectly, manual alignment of the histology images may be necessary.

**Fig. 14.**
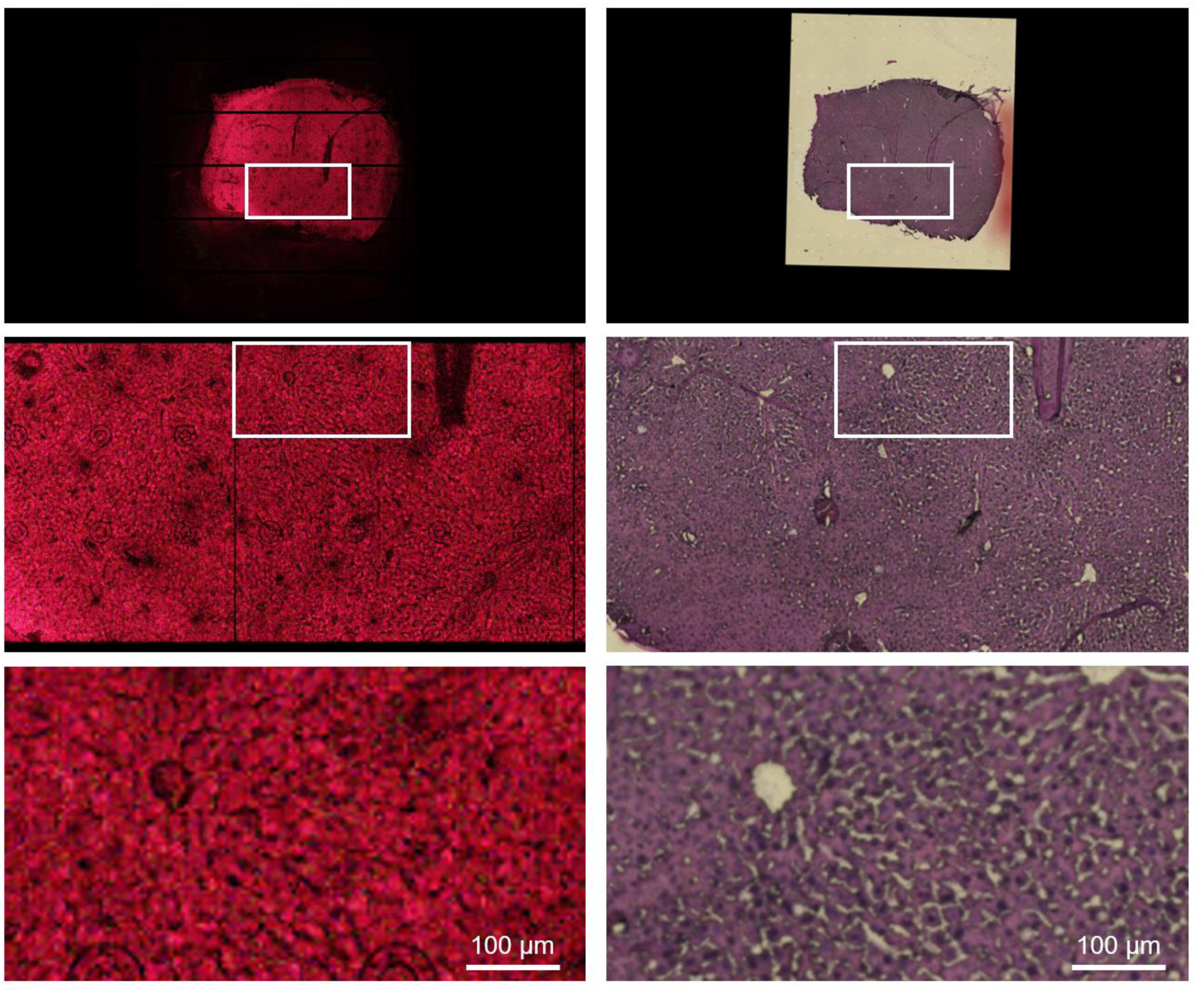
Comparison between sge (left) and aligned histology (right) images. Boxed areas are sequentially magnified below. The aligned histology image is the output image from step 2-3), a tiff file with histology imaging information that has the same dimension as in smatch (Fig. 12) and sge (Fig. 13) images.

#### 2-4) (Optional) All-in-one analysis from STEP 1-4) to 2-3)

1. All the steps described steps 1-4) and 2-3) can be performed all at once using the *all* step in the NovaScope pipeline. Following the instruction, the user needs to prepare the configuration file in YAML format, which should contain the locations of input FASTQ files, parameter settings, and the locations of the reference genome index.

### STEP 3. Exemplary Downstream Analysis

#### 3-1) Data Preprocessing

1. Convert the SGE matrix to FICTURE format, where each row contains, X/Y coordinates, gene names, identifiers, and observed counts.
2. Using make_spatial_minibatch.py script in FICTURE, reformat the input file by assigning minibatch label, and by reordering the data so that they are locally contiguous.

#### 3-2) Unsupervised Inference of Cell Type Factors using Latent Dirichlet Allocation

1. Using make_dge_univ.py script in FICTURE, given a specified size of hexagons, segment the raw SGE matrix into hexagons and store them into a spatial gene expression (SGE) matrix in the format compatible with FICTURE.
2. Using the lda_univ.py script in FICTURE, perform an unsupervised learning of cell type factors using Latent Dirichlet Allocation (LDA).
3. Examine the output image of overlapping hexagonal factors (Fig. 15) to see if the inference of cell type factors is reasonable (Fig. 16).

#### 3-3) (Optional) Use External Tools to Infer Cell Type Factors

1. Using make_sge_by_hexagon.py script in FICTURE, with a specified size of hexagons, segment the raw SGE matrix into hexagons and store them into a SGE matrix consistent to 10x Genomics format, which can be loaded in other single-cell or spatial genomics software tools such as Seurat^28^ and Scanpy^30^. In this format, the barcodes contain the spatial coordinates of each hexagon instead of nucleotide sequences.
2. Using external software tools such as Seurat, identify cell type clusters for each hexagon using the hexagonal DGE matrix as input. Examine whether the inferred clusters make sense.
3. Using the external_cluster_to_count_matrix.py, generate the model matrix for individual cell types in the FICTURE format for pixel-level decoding. Other cell type factors derived from single-cell reference data or cell segmentation can also be converted into the model matrix for pixel-level decoding.

#### 3-4) Pixel-level decoding of cell type factors using FICTURE

1. Using the transform_univ.py script in FICTURE, group the pixel-level data from the SGE matrix in FICTURE format from 3-1) into hexagons, then transform these hexagons into factor space based on the input model matrix from 3-2) or 3-3). The output files include a tab-delimited file that provides a spatial summarization of gene expressions by hexagons, weighted by the hexagon’s factor probabilities.
2. Using the slda_decode.py script in FICTURE, perform pixel-level decoding of the model matrix on individual pixels. The output file is a tab-delimited file containing the posterior count of factors on individual pixels.
3. Using the de_bulk.py script in FICTURE, identify marker genes that are associated with each factor.
4. Using the factor_report.py script in FICTURE, create an HTML file summarizing individual factors and marker genes.
5. Using the plot_pixel_full.py script in FICTURE, create a high-resolution image of cell type factors for individual pixels. The color of each factor is consistent with the HTML report above.
6. (Optional) Using the plot_pixel_single.py script in FICTURE, create high-resolution images for individual factors separately.
7. Examine the output image of pixel-level decoding (Fig. 18, 20) to see if the updated inference of cell type factors (Fig. 19) is reasonable.

## TROUBLESHOOTING

### A. Experimental Procedures

Step: 1-2

Problem: Abnormal dicing patterns; the chip does not break in a clean line.

Possible Reason: Improper settings for the Precision Glass Cutter.

Solution: Adjust the glass knife tightness according to the Precision Glass Cutter manual. Incorrect depth of scoring could lead to abnormal dicing patterns.

Step: 2-5

Problem: Freezing artifact in histology images.

Possible Reason: Most of these artifacts are caused by problems during tissue freezing.

Solution: Use only tissues properly frozen according to Step 2-1. Do not leave the cryomold in isopentane for an extended period. Adjust the freezing time in isopentane so that freezing damage is limited to a minimum.

Step: 3-5

Problem: PCR product concentration is too low.

Possible Reason: Failure in array processing, RNA capture, or reverse transcription. Suboptimal tissue permeabilization. Tissue RNA degradation or low RNA concentration in the tissue. Low occupancy of tissue on the chip.

Solution: Ensure adequate humidity is provided in the humidified chamber. Ensure that evaporation is minimal during long-term incubation of solutions. Ensure that tissues are well permeabilized (see Step 3-1 for details). Ensure that tissue RNA quantity and quality are high (see Step 2-2 for QC procedures). Ensure that the tissue overlaid on the chip covers a substantial area, which is crucial for obtaining enough material for library construction.

Step: 3-7

Problem: Small-sized product is contaminating the index library.

Possible Reason: Insufficient size selection.

Solution: Size selection, either through gel elution or bead-based purification, should be repeated to completely remove the undesired low molecular weight product.

Step: 3-7

Problem: Low index PCR product concentrations.

Possible Reason: Low template concentrations.

Solution: Increase template input. Alternatively, increase the reaction volume of index PCR (in multiple PCR tubes) or extend PCR cycles so that adequate library quantity is obtained for sequencing.

**Fig. 15.**
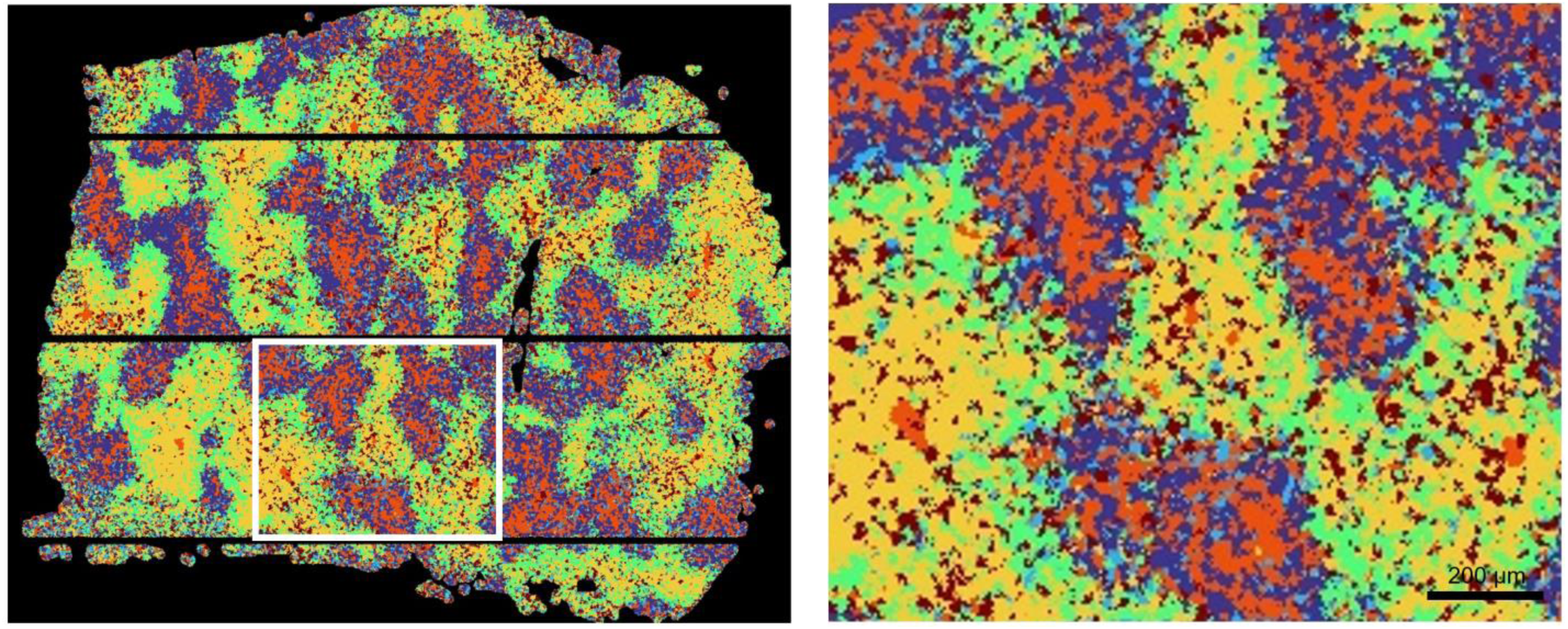
An exemplary output image from step 3-2, illustrating distinct zonation of hepatocellular factors. Refer to Fig. 16 for the color-coding scheme. The LDA settings are d18 (10 µm-sided hexagons) and nF6 (number of factors = 6). Boxed area is magnified on the right.

**Fig. 16.**
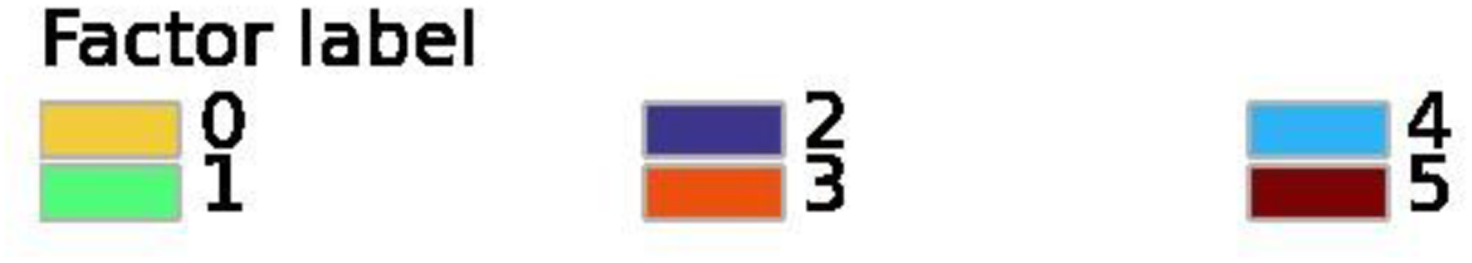
Factor color legend for Fig. 15. Factors 0-4 represents metabolically zonated hepatocytes along the portal-central axis. 0: Extreme periportal, 1: Periportal, 2, 4: Central, 3: Extreme Central. 5: Red blood cells.

**Fig. 17.**
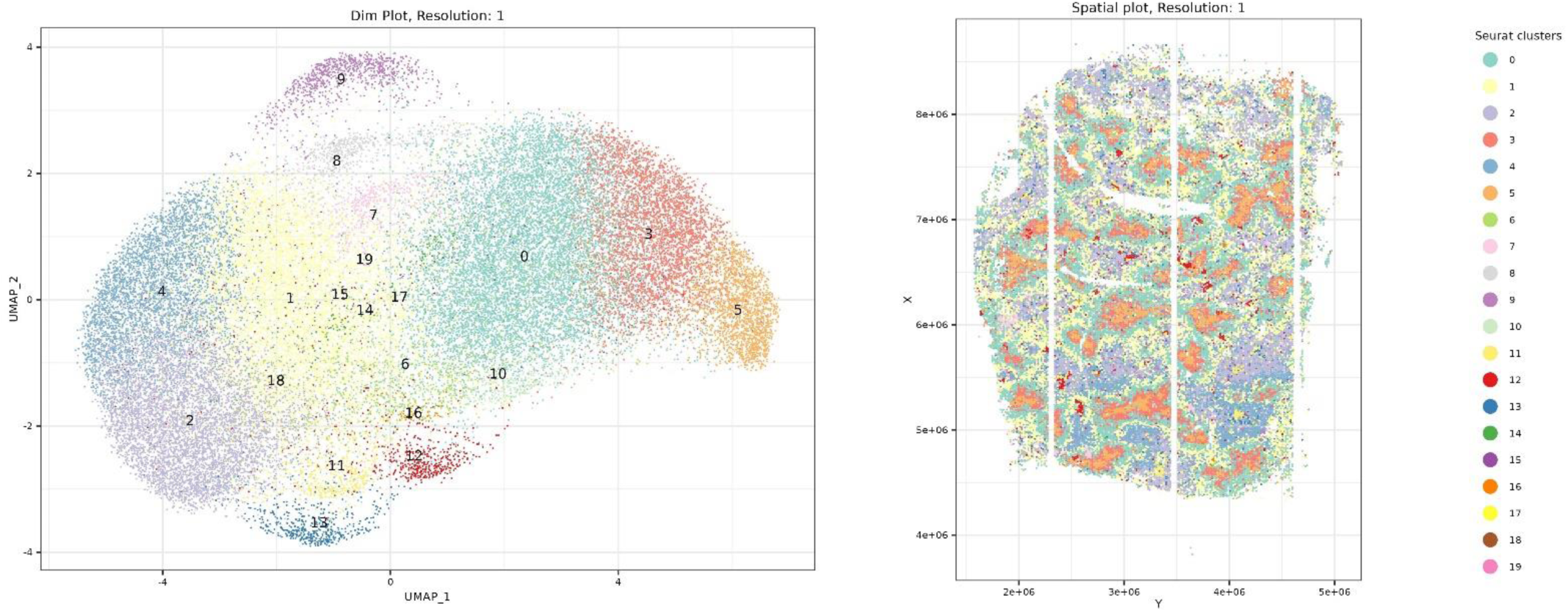
An exemplary multi-dimensional clustering result obtained using Seurat. (A) UMAP manifold visualizing 20 clusters representing different cell types. Notable clusters include 0, 3, 5: Pericentral hepatocytes, 2, 4: Periportal hepatocytes, 1: Midzone hepatocytes, 6, 16: B cells, 7: Cells undergoing antiviral response, 9: Red blood cells, 10: hepatic stellate cells, 11: Myofibroblasts and macrophages, 12: Hepatocytes with damage response, 13: Cholangiocytes/liver progenitors. The clusters were obtained using following parameters: 10 µm-sided hexagons (d18) and FindCluster resolution of 1.00. These results reproduce the liver results that were described in the original Seq-Scope paper^9^.

**Fig. 18.**
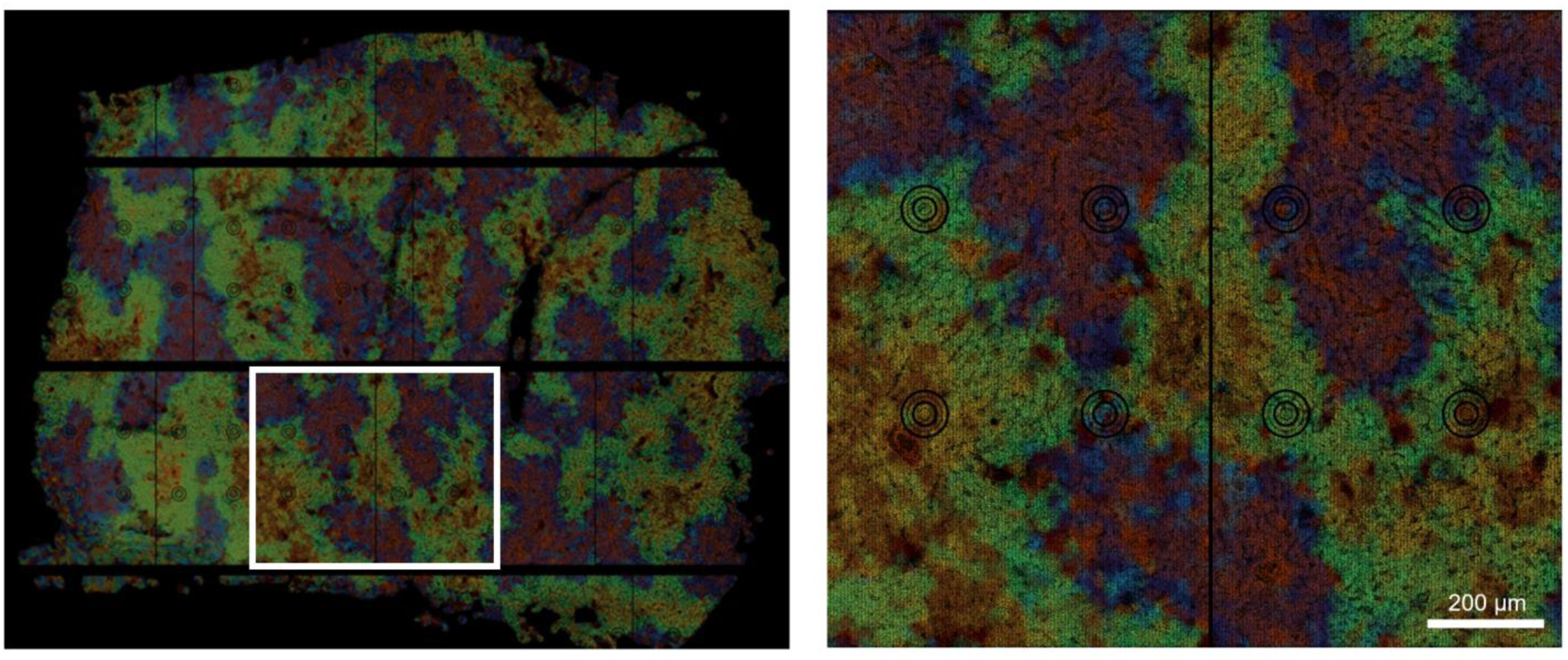
An exemplary pixel-level output image from step 3-4) showing clear zonation of hepatocellular factors. The color-coding scheme is the same as shown in Fig. 16. Boxed area is magnified on the right.

**Fig. 19.**
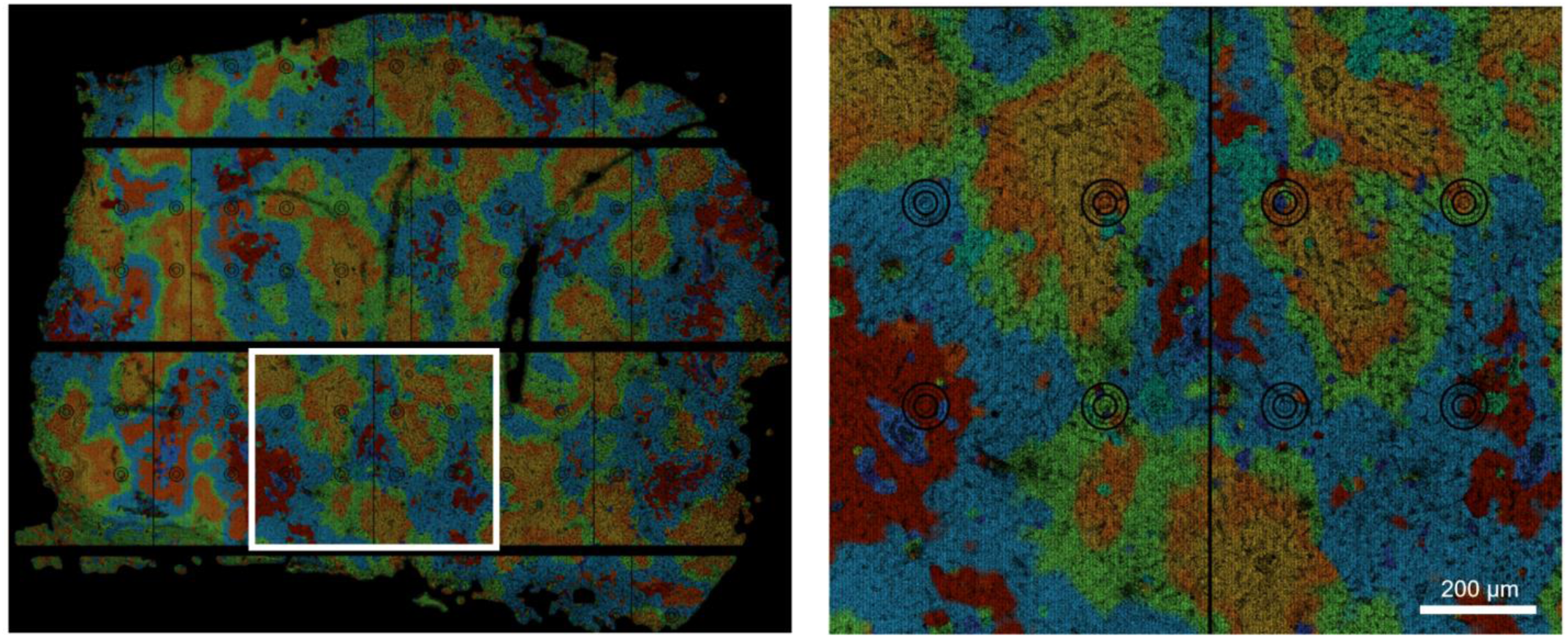
Pixel-level projection of Seurat cluster factors learned from the analysis shown in Fig. 17. The color coding scheme for each factor is presented in Fig. 20. Boxed area is magnified on the right.

### B. Computational Procedures

Step: 1-3

Problem: The proportion of non-unique spatial barcode is high (>20%), or the total number of unique barcodes per tile is low (< 2 million).

Possible Reason: Underclustering, overclustering, or equipment malfunction.

Solution: Check the cluster occupancy using the Illumina Sequence Analysis Viewer (SAV) and adjust the library concentration as needed. Consult with the Illumina service team to determine if there are mechanical problems in the system affecting the sequencing runs.

Step: 2-1

Problem: The output image of the spatial distribution of matching barcodes does not match to the tissue area. The tissue shape pattern is unseen or diffuse from the spatial dataset, or shows as an inverse image (tissue region does not reveal the transcriptome while the area outside the tissue reveals many transcripts).

Possible Reason: Improper tissue permeabilization.

Solution: Optimize the tissue permeabilization condition using the method described in the original spatial transcriptomics paper^15^ or the tissue optimization kit commercially available by 10x Visium^16^. Alternatively, optimize the permeabilization condition directly within the Seq-Scope workflow, using different imaging areas of the Chip. Adjust the tissue section thickness (e.g., from 10 µm to 5 µm) if necessary.

Step: 2-2

Problem: Unspliced/genomic reads are discovered from the tissue, while spliced reads are lacking. Possible Reason: RNA is degraded, so the chip is capturing genomic DNA instead.

Solution: Ensure that tissue RNA quality is intact by running the STEP A-2-2 RNA QC procedure.

Step: 2-2 and afterwards

Problem: Transcriptome density is too low, so cell type mapping is not properly conducted. Possible Reason: RNA degradation, library preparation errors, tissue permeabilization issues.

Solution: As discussed above, check tissue RNA quality, ensure reaction solutions have not dried out, and ensure the tissue permeabilization condition is optimal.

## TIMING

### A. Experimental Procedures

STEP 1. Seq-Scope Chip Production (1^st^-Seq): ∼33 hours per flow cell (≥ 72 experiments in parallel)

1-1) NovaSeq 6000 Operation: ∼15 hours (could include overnight sequencing run)

1-2) Flow Cell Disassembly and Dicing: ∼2 hours

1-3) Seq-Scope Chip Surface Treatment: ∼16 hours (including one overnight step)

STEP 2. Tissue Preparation, Attachment, and Imaging: ∼3 hours per Chip (≥ 4 experiments in parallel)

2-1) [Not included in the overall TIMING] Tissue Preparation: ∼1 hour

2-2) Tissue sectioning and attachment: ∼2 hours

2-3) [Not included in the overall TIMING] Quality check for tissue histology and RNA integrity: ∼2 hours 2-4) Tissue Fixation: 10 minutes

2-5) Tissue H&E Staining and Imaging: ∼1 hour

STEP 3. Library Construction and Sequencing (2^nd^-Seq): ∼20 hours plus sequencing time (≥ 4 experiments in parallel)

3-1) mRNA Release and Reverse Transcription: ∼16 hours (including one overnight step)

3-2) Tissue Removal: 1 hour 30 minutes

3-3) Secondary Strand Synthesis: 2 hours 30 minutes

3-4) Purification of the Secondary Strand Solution: ∼30 minutes

3-5) Library Amplification PCR Reaction and Purification: ∼3 hours

3-6) Indexing PCR and Size Selection: ∼2 hours

3-7) Library Sequencing: Varies depending on the workflow, platform, scale, and pooling. In our workflow involving two sequencing runs (MiSeq-based SequencingQC and NovaSeq-X-based final sequencing), this step requires ∼27 hours including two overnight runs.

### B. Bioinformatic Procedures

These estimations are based on one full NovaSeq 6000 S4 flow cell (1st-Seq) and a 5GB FASTQ file with 200M reads (2nd-Seq). The system utilized 2–12 CPUs and 14–84GB of RAM, depending on the tasks. Generally, runtime depends on the FASTQ size and the computational capacity.

STEP 1. Generation of Spatial Barcode Maps

1-1) Install NovaScope pipeline: 10 minutes

1-2) (Optional) Generate 1^st^-seq FASTQs from BCL files: ∼3 hours

1-3) Build a spatial barcode map for individual tiles: ∼3 hours

1-4) Build a spatial barcode map for each Chip: 20 minutes

STEP 2. Sequence Alignment and Spatial Gene Expression (SGE) Matrix Generation.

2-1) Match 2^nd^-Seq reads to spatial barcodes of the corresponding Chip: ∼10 minutes

2-2) Align the 2^nd^-Seq reads to the reference genome and generate an SGE matrix: ∼30 minutes

2-3) (Optional) Histology alignment based on fiducial mark and tissue density: ∼20 minutes

2-4) (Optional) All-in-one analysis for STEP 1-4) to 2-3): 1.5 hours

STEP 3. Exemplary Downstream Analysis

3-1) Data Preprocessing and Generation of Hexagonal SGE: 10 minutes

3-2) Unsupervised Inference of Cell Type Factors using Latent Dirichlet Allocation: 10 minutes

3-3) (Optional) Use External Tools to Infer Cell Type Factors: ∼10 minutes.

3-4) Pixel-level decoding of cell type factors using FICTURE: 15 minutes.

## PAUSE POINTS

1-1) PAUSE POINT: The sequenced flow cell can be stored at 4°C for up to 2 months. 1-2) PAUSE POINT: The diced chip can be stored at 4°C for up to 2 months.

1-3) PAUSE POINT: The surface-treated chip can be stored at 4°C for up to 1 week. 2-1) PAUSE POINT: The frozen sample can be stored at –80°C for up to 6 months.

2-5) PAUSE POINT: The H&E-stained chip, after an 80% ethanol wash and drying, can be stored at 4°C for up to 1 week.

3-4) PAUSE POINT: The purified secondary strand can be stored at –20°C for up to 6 months.

3-5) PAUSE POINT: The purified Library Amplification PCR product can be stored at –20°C for up to 6 months. 3-6) PAUSE POINT: The final library can be stored at –20°C for up to 6 months.

## ANTICIPATED RESULTS

We map HDMI in 60 tiles (10 × 6) arranged specifically for each imaging area (Fig. 11); typically, a single imaging area would occupy 30–36 (5 × 6 or 6 × 6) tiles that are located at the center of the 60-tile area (Fig. 12). Approximately 50% of all 2^nd^-Seq reads would be mapped to 1^st^-Seq tile areas (Fig. 12). The rest of the reads are unmapped due to various sources, such as sequencing errors, duplicate HDMI barcodes that we remove from the dataset, HDMI from clusters that fail to sequence, or HDMI clusters outside of the sequenced areas. From these 2^nd^-Seq reads whose HDMI was mapped to 1^st^-Seq tile areas, approximately 50% are mapped uniquely to genic regions (Fig. 13). Other reads may be mapped to ribosomal sequences, mapped to multiple regions (conserved sequences), mapped to intergenic regions (DNA capture as described in the Troubleshooting section), or unmapped to genome (poly-A-only capture). Transcript density varies extensively across different tissues; however, an average transcript density of 1–2 UMI/µm^2^ is typically necessary to successfully map different cell types at the cellular level.

In this protocol article, we demonstrated spatial transcriptome profiling of mouse liver tissue, reproducing all major findings in our original Seq-Scope publication^9^ with slightly better transcriptome capture efficiency (currently 8.7 UMI/µm^2^ over the previous 6.7 UMI/µm²) and a substantially larger continuous imaging areas (currently 7 mm × 7 mm over our previous 1 mm-diameter circles). In this experiment, tissues covered approximately 12 mm^2^ of the space. A total of 2.61 billion 2^nd^-Seq reads were analyzed and 1.32 billion reads were mapped to the corresponding imaging area (60 tiles with 6 swaths and 10 lengths). Among these, 54% of the reads were uniquely mapped to genes and most of these were spliced RNAs, while some unspliced RNAs were detected. As a result, the dataset revealed 21.9 million pixels and 108.8 million transcripts. The dataset is characterized by very high resolution, preserving microscopic details of cellular and subcellular structures of hepatocytes consistent with underlying histology (Fig. 14).

The dataset is analyzed through two exemplary methods utilizing FICTURE, which enables pixel-level inference of spatial factors with raw resolution details^29^. The first analysis utilizes LDA-based factor learning, which is a default option for FICTURE analysis; the second analysis utilizes multi-dimensional clustering algorithm implemented in Seurat package^28^.

Both methods proved effective in identifying metabolically zonated hepatocytes and were also successful in recognizing various cellular states, including hepatocytes undergoing damage or antiviral responses, as well as non-hepatocytes like hepatic stellate cells, macrophages/myofibroblasts, cholangiocytes, and red blood cells (Fig. 15-17). FICTURE effectively projected these findings onto the histological space with pixel-level resolution (Fig. 18-20).

**Fig. 20.**
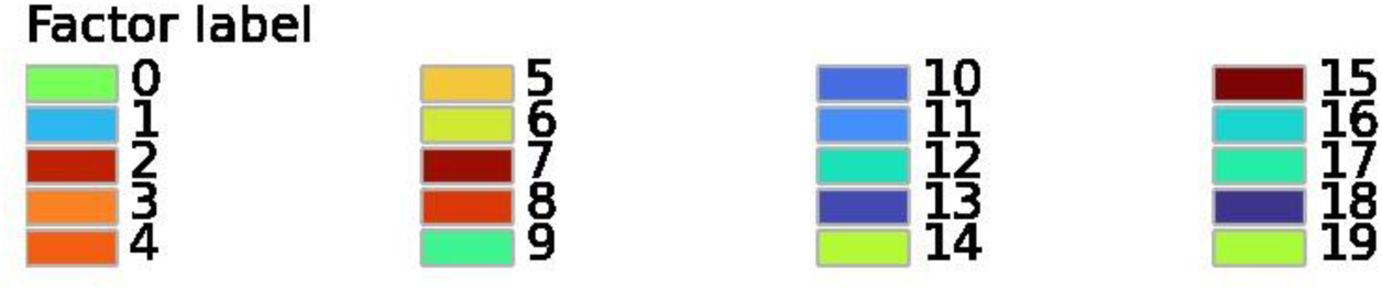
Factor color legend for Fig. 19. Color code for each factor whose number is corresponding to what was presented in Fig. 17.

The dataset described above was obtained from deep sequencing (2.61 billion reads) of the 2nd-Seq library. Importantly, such deep sequencing may not be absolutely required because useful levels of information could be attainable with relatively shallow 2^nd^-Seq library sequencing (∼163 million reads), which only requires a low-cost budget (∼1.5% of shared NovaSeq-X flow cell sequencing, which costs around $250). In this case, the total cost of running Seq-Scope (including array sequencing and library sequencing) could be lower than $500. The shallower sequencing revealed 10.9 million pixels and 18.6 million transcripts, leading to a final transcriptome density of 1.5 UMI/µm^2^, which is smaller than the deeper sequencing (8.7 UMI/µm^2^). Nevertheless, the data still are able to identify major cell types pertaining to liver cell diversity, and also showed localized gene expression across the portal-central metabolic zonation axis (Fig. 21), as originally demonstrated by our earlier study^9^. Therefore, Seq-Scope can offer low-cost high-resolution spatial transcriptomics analysis, while higher-cost deep sequencing can be used to reveal more comprehensive information based on the user’s preference.

**Fig. 21.**
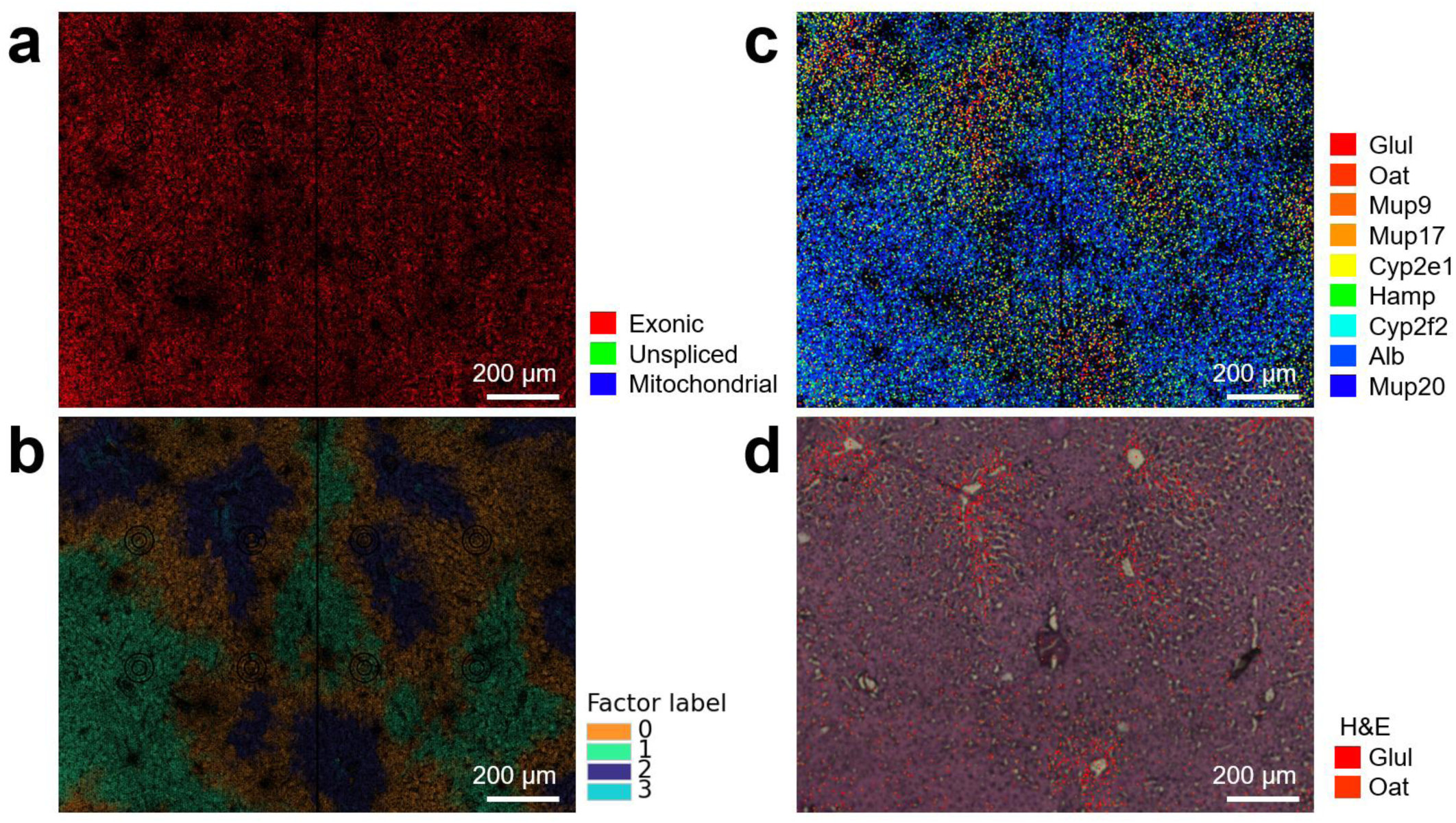
Seq-Scope analysis using shallowly sequenced liver data (∼163 million 2^nd^-Seq read inputs). (a) sge image produced as in Fig. 13. (b) FICTURE projection image produced as in Fig. 18 with d24 (14 µm-sided hexagons) of shallowly sequenced data. (c) Spatial gene expression plot visualizing each indicated transcript with dots of corresponding color and varying alpha levels. Warm colors represent pericentral expression while cold colors represent periportal expression. Abundantly expressed genes, such as Alb, were plotted with lower alpha levels. (d) H&E histology overlaid with spatial gene expression plots visualizing central area-specific *Glul* and *Oat* transcripts.

## SUPPLEMENTARY INFORMATION

**Supplementary Data 1.** 3D model (STL) file for the Custom Frame Adapter described in the protocol. The STL file can be used to fabricate the adapter in most 3D printing service centers. We printed the adapter on a Stratasys J850 using the default white material.

**Supplementary Data 2.** Sketch drawing and specification of the Custom Silicone Isolator (Grace Bio-Labs, cat. no. JTR25-A-1.0, RD501346). Information in this PDF is sufficient to reproduce the part with the same specifications applied by Grace Bio.

**Supplementary Video 1:** NovaSeq 6000 S4 Flow Cell Disassembly. The scalpel is used to separate the flow cell into its three main components. This video demonstrates the entire procedure of the flow cell disassembly. During the demonstration, viewers may notice that a minor crack was introduced on the top layer of the flow cell. This layer is thin and, therefore, prone to breakage. However, as long as the crack is outside the imaging area described in Fig. 3d (B02–B10 and T02–T10), it does not interfere with subsequent procedures. Furthermore, even if the crack damages some imaging areas, other intact areas can still be effectively utilized without any issues.

**Supplementary Video 2.** Seq-Scope Chip Dicing Procedure. The Chip was diced from the disassembled top and bottom layers of a NovaSeq 6000 S4 flow cell using the Precision Glass cutter.

## AUTHOR CONTRIBUTION STATEMENTS

YK, CSC, AP, MS, JEH, MK and JHL developed the experimental part of the protocol. WC, YH, YS, JX and HMK developed the computational part of the protocol. EP, OIK, TW, HMK and JHL developed the sequencing part of the protocol. YK, WC, CSC, HMK and JHL prepared a draft. All authors revised, reviewed, and approved the final version.

## ORCID FOR CORRESPONDING AUTHORS

Jun Hee Lee, 0000-0002-2200-6011

Hyun Min Kang, 0000-0002-3631-3979

## ACKNOWLEDGEMENTS

The work was supported by the Taubman Institute Innovation Project (to HMK and JHL), the NIH (T32AG000114 to YK and CSC, K01AG061236 to MK, R01AG079163 to MK and JHL, R01HG011031 and HHSN268201800002I to HMK, and R01DK133448 and UH3CA268091 to JHL), Technology Transfer Talent Network (T3N) Postdoctoral Fellowship (to YK), and Gleen Foundation Core grant (to JHL).

