## Supplementary figures and images for "Seq-Scope Protocol: Repurposing Illumina Sequencing Flow Cells for High-Resolution Spatial Transcriptomics"

### Supplementary Data 2.pdf

#### Silicone Isolator

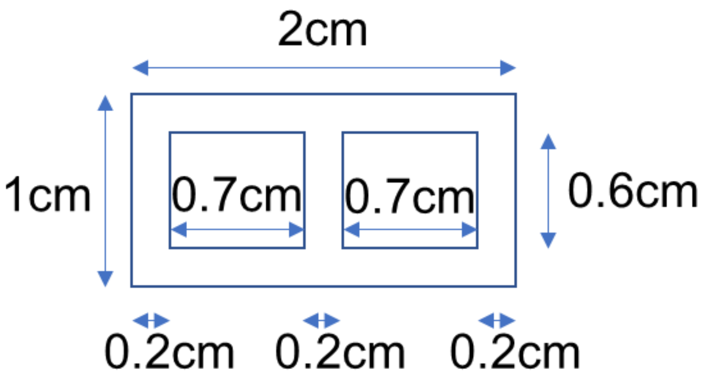


Color: Red
Adhesive: One side
Chamber Depth: 1mm
Part Count: 500
